# Evolutionary trade-offs between unicellularity and multicellularity in budding yeast

**DOI:** 10.1101/347609

**Authors:** Jennie J. Kuzdzal-Fick, Lin Chen, Gábor Balázsi

## Abstract

Multicellular organisms appeared on Earth through several independent major evolutionary transitions. Are such transitions reversible? Addressing this fundamental question entails understanding the benefits and costs of multicellularity versus unicellularity. For example, some wild yeast strains form multicellular clumps, which might be beneficial in stressful conditions, but this has been untested. Here we show that unicellular yeast evolves from clump-forming ancestors by propagating samples from suspension after larger clumps have settled. Unicellular yeast strains differed from their clumping ancestors mainly by mutations in the *AMN1* (Antagonist of Mitotic exit Network) gene. Ancestral yeast clumps were more resistant to freeze/thaw, hydrogen peroxide, and ethanol stressors than their unicellular counterparts, while unicellularity was advantageous without stress. These findings inform mathematical models, jointly suggesting a trade-off between the benefits and downsides of multicellularity, causing bet-hedging by regulated phenotype switching as a survival strategy in unexpected stress.

## 1. INTRODUCTION

Multicellularity has evolved over 20 different times on Earth, leading to complex life forms in algae, animals, plants, and fungi (Grosberg and Strathmann 2007). What forces contributed to the emergence and maintenance of multicellularity, and could reversals to unicellularity occur? Over 100 years ago Louis Dollo hypothesized that phenotypic effects of evolutionary processes must be irreversible (Dollo 1893), but this has since become controversial (Collin and Miglietta 2008). Possible exceptions include the loss of flight in winged dinosaurs and birds (Paul 2002), of parasitic traits in dust mites (Klimov and OConnor 2013), or of protein sequence changes (Soylemez and Kondrashov 2012; McCandlish et al. 2016). How strictly Dollo’s Law applies to major evolutionary transitions (Maynard Smith and Szathmáry 1998), such as the relatively easy transition to multicellularity (Grosberg and Strathmann 2007; Ratcliff et al. 2012) is insufficiently explored. Addressing this problem requires charting the environment dependent downsides and benefits of unicellularity versus multicellularity. For example, multicellularity benefits dispersal in sparse nutrient conditions (Kuzdzal-Fick et al. 2007; Smith et al. 2014), stress resistance (Smukalla et al. 2008), nutrient acquisition (Koschwanez et al. 2011) and predator protection (Pentz et al. 2015). Implicitly, unicellularity is disadvantageous in such conditions – yet it improves growth without stress (Smukalla et al. 2008). We set out to study these trade-offs in budding yeast.

Contrary to the classical definition of yeasts as single-celled fungi, some *Saccharomyces cerevisiae* strains exhibit multicellular phenotypes (Reynolds and Fink 2001; Cap et al. 2012; Andersen et al. 2014), such as aggregation into flocs (Smukalla et al. 2008), flors (Zara et al. 2009) and pattern formation on agar plates (Reynolds and Fink 2001; Kuthan et al. 2003; Chen et al. 2014). Other yeast strains form multicellular “clumps” that differ from flocs and flors in their mechanism of formation and underlying genetics (Li et al. 2013). Instead of cell aggregation, clumps form by incomplete daughter cell separation, as budding continues, but daughter cells remain attached to the mother cell (Kuranda and Robbins 1991). Such yeast clumps aid nutrient acquisition in sucrose (Koschwanez et al. 2011), but their role in stress is unclear.

Understanding the costs and benefits of social traits in yeast could elucidate general forces that maintain or convert unicellularity to multicellularity (Maynard Smith and Szathmáry 1998) and back. The existence of unicellular and clumpy yeast in nature (Wloch-Salamon et al. 2013) suggests condition-dependent benefits and downsides, and bidirectional transitions between unicellularity and multicellularity. Could clumps provide protection from environmental stress as flocs do (Smukalla et al. 2008) while being disadvantageous in normal conditions? More broadly, could reverse transitions to unicellularity occur and what are the evolutionary forces that aid or prevent such reverse transitions?

To address these questions, here we compared how various environmental stressors affect the growth of genetically similar clump-forming and unicellular “EvoTop” yeast cells that we obtained by reversing the strategy of “snowflake” yeast evolution (Ratcliff et al. 2012). Sequencing and comparing the genomes of the clumping ancestor and single-celled “EvoTop” lines revealed unique missense and nonsense mutations in the *AMN1* gene, which is associated with multicellularity (Yvert et al. 2003; Li et al. 2013). Clump-forming ancestral cell lines grew faster relative to untreated controls than EvoTop lines after exposure to rapid freeze/thaw, 1% ethanol, and 150 μM H_2_O_2_ stressors, indicating that clumping provides resistance to chemical and physical stresses. On the other hand, clumping hampered growth in the absence of stress, suggesting a trade-off between the benefits and downsides of multicellularity versus unicellularity. Mathematical models captured these observations and revealed how multicellularity causes regulated phenotype switching that aids survival during unexpected periods of stress. Overall, this work sheds light on the genetic bases, as well as on the costs and benefits, of unicellularity versus clumping multicellularity in yeast, with implications for bidirectional transitions between other unicellular and multicellular life forms.

## 2. MATERIALS AND METHODS

### Yeast strains

We used three strains of the budding yeast *S. cerevisiae* in this study. The first one was TBR1 (∑1278b strain 10560-23C; MATα, ura3-52, his3::hisG, leu2::hisG), a segregant obtained by multiple crosses of baking strains “Yeast Foam” and 1422-11D that carries 3.2 single-nucleotide polymorphisms per kilobase compared to the standard laboratory strain S288c (28). The second was the standard laboratory strain BY4742 (S288C-derivative, MATα his3Δ1 leu2Δ0 lys2Δ0 ura3Δ0). The third one was KV38 (haploid strain obtained from the wild strain EM93, the ancestor of S288c and source for 90% of its gene pool).

### Selection for unicellular yeast

To initiate 3 replicate lines of the haploid yeast strain TBR1, we inoculated 3 tubes of 2 ml yeast peptone dextrose (YPD) medium (10 g yeast extract, 20 g bacto peptone, 2% glucose per L) with single TBR1 colonies. We allowed each culture to grow overnight, froze aliquots of these “ancestral” cultures (TBR1 A, B, & C), and then prepared two 100x dilutions of each culture in 2 ml YPD to obtain matched pairs of TBR1 A, B, and C for starting the EvoTop and EvoControl treatment lines. Each line reached stationary phase by growing for 24 hours in a 30°C shaking LabNet 311DS incubator at 300 rpm. After removal from the shaking incubator, large clumps should settle faster than single-cells and small clumps. For the EvoTop treatment lines, we vortexed each tube before allowing them to remain in a 30°C MyTemp Mini Digital static incubator for 45 minutes before taking a 20 μl sample from the top of the liquid culture to inoculate a new tube with 2 ml of YPD for growth overnight under the previously described conditions. We performed the selection procedure each day over 4 weeks, for a total of 28 rounds of selection. Over the course of the selection experiment, we maintained parallel EvoControl lines by vortexing each strain before selecting 20 μl samples for each transfer into new tubes with 2 ml YPD. We froze ancestral samples at the beginning of the experiment and samples of each EvoTop and EvoControl line (700 μl cell solution with 300 μl 80% glycerol) in a −80°C freezer every 3 to 4 days, including the final cultures. We followed the same protocol with another haploid clump-forming strain, KV38 (Smukalla et al. 2008), and with the haploid unicellular laboratory strain BY4742, a S288c derivative (Brachmann et al. 1998), as a control.

### Estimating cell and clump sizes

To determine cell size and clump size of each EvoTop, EvoControl, and ancestral line, we used a Nexcelom Auto M10 automated cell counter to analyze 10x dilutions in YPD of overnight cultures (grown in YPD in a 300 rpm shaking 30°C LabNet 311DS incubator) started from frozen samples of each line from the ancestral state through to the end product. For cell size, we set the Cellometer Auto cell type parameters as follows: cell diameter minimum of 2.0 μm and maximum of 9.0 μm, roundness of 0.10, and contrast enhancement of 0.40, with a decluster edge factor of 0.5 and Th factor of 1.0. We measured clumps with the following parameters: cell diameter minimum of 2.0 μm and maximum of 40.0 μm, roundness of 0.10, and contrast enhancement of 0.40, with “Do not Decluster Clumps” selected. We increased the maximum cell diameter of clumps to 100 μm for samples where the program indicated that some clumps were larger than 40 μm in diameter. Diameters from 10x dilutions of live samples were also measured in this manner a number of times towards the beginning and end of the selection experiment.

We combined clump and cell diameter data gathered during the experiment with data from the lines taken out of the freezer and then averaged together all clump or cell data points by line (within strain and treatment) for days that had both, including the initial “Ancestor” and final “EvoTop” and “EvoControl” days. We used the *userfriendlyscience* package in R version 3.4.4 (R_Core_Team 2013; Peters 2018) to perform a one-way ANOVA analysing the effect of treatment on clump diameter or the effect of treatment on cell diameter for TBR1, BY4742, and KV38, followed by Games-Howell post-hoc tests.

### TBR1 sequencing analysis

The DNA from the ancestor and each of the three TBR1 EvoTop isolates was extracted from cultures started from individual colonies that were anticipated to be clonal. Sequencing analysis was performed as described in the Supplementary Material.

### Assays of stress resistance

To compare the differences in stress response between the clump-forming TBR1 ancestor and its “single-celled” descendants, we isolated colonies from each ancestral and EvoTop TBR1 line (A, B, and C) that were good representatives of these phenotypes. We diluted exponentially growing cells started from individual colonies and placed 1.5 ml of each dilution into 2 separate 2 ml microcentrifuge tubes to start a control and experimental stress treatment. Stress resistance was tested as described in the Supplementary Material. We used SAS Studio 3.6 to run a N-Way full factorial ANOVA (procedure GLM) and Tukey post-hoc tests to analyze strain, line, and stress effects on the relative growth of TBR1 and BY4742 cells.

### TBR1 BY4742 control growth rates

We ran a N-Way ANOVA (procedure GLM, Type I SS) in SAS Studio 3.6 to analyze the effects of strain, phenotype, and line and strain X phenotype and phenotype X line interactions on the growth (proportion change) of unstressed TBR1 and BY4742 clumps and single-cells. For this analysis, TBR1 ancestors and BY4742 EvoTops were considered “clumps,” while TBR1 EvoTops and BY4742 ancestors were considered “single-cells.”

### Data sharing

Whole-genome sequence data is available from NCBI assembled at BioProject PRJNA388338.

Additional data is available at http://www.openwetware.org/wiki/Image:ClumpingData-Scripts.zip

## 3. RESULTS

### The TBR1 yeast strain shows the clumping phenotype

To explore the bidirectional transitions and fitness effects between unicellularity versus multicellularity, we focused on the haploid *S. cerevisiae* strain TBR1 (see the Methods), a segregant obtained by multiple crosses of baking strains that carries thousands of polymorphisms relative to the standard laboratory strain S288c (Dowell et al. 2010) and develops wrinkly patterns on soft agar plates (Reynolds and Fink 2001; Chen et al. 2014). We asked whether TBR1 cells would also be capable of clump formation by incomplete separation. Thus, we compared phenotypes related to clump formation in the TBR1 strain and the standard laboratory strain BY4742, a haploid-derivative of S288c.

Strains capable of clump formation tend to settle to the bottom of the culture tube over time. Therefore, we first visually tested the settling of these two yeast strains 45 minutes after removal from the shaking incubator. We noticed that the entire liquid culture medium was still uniformly turbid for the laboratory strain BY4742. In contrast, the TBR1 strain formed a vertical gradient of turbidity, increasing from top to bottom (Figure 1A). Next, we examined by microscopy how these different settling behaviours correlated with sizes and shapes at the cellular level. The standard laboratory strain appeared mainly as single-cells, doublets and occasional triplets (Figure 1B, top), whereas the TBR1 strain appeared as small, but compact and regular-shaped clumps and multiplets (Figure 1B, bottom), suggesting that clumps must split or must shed cells to limit their sizes.

**Figsure 1.**
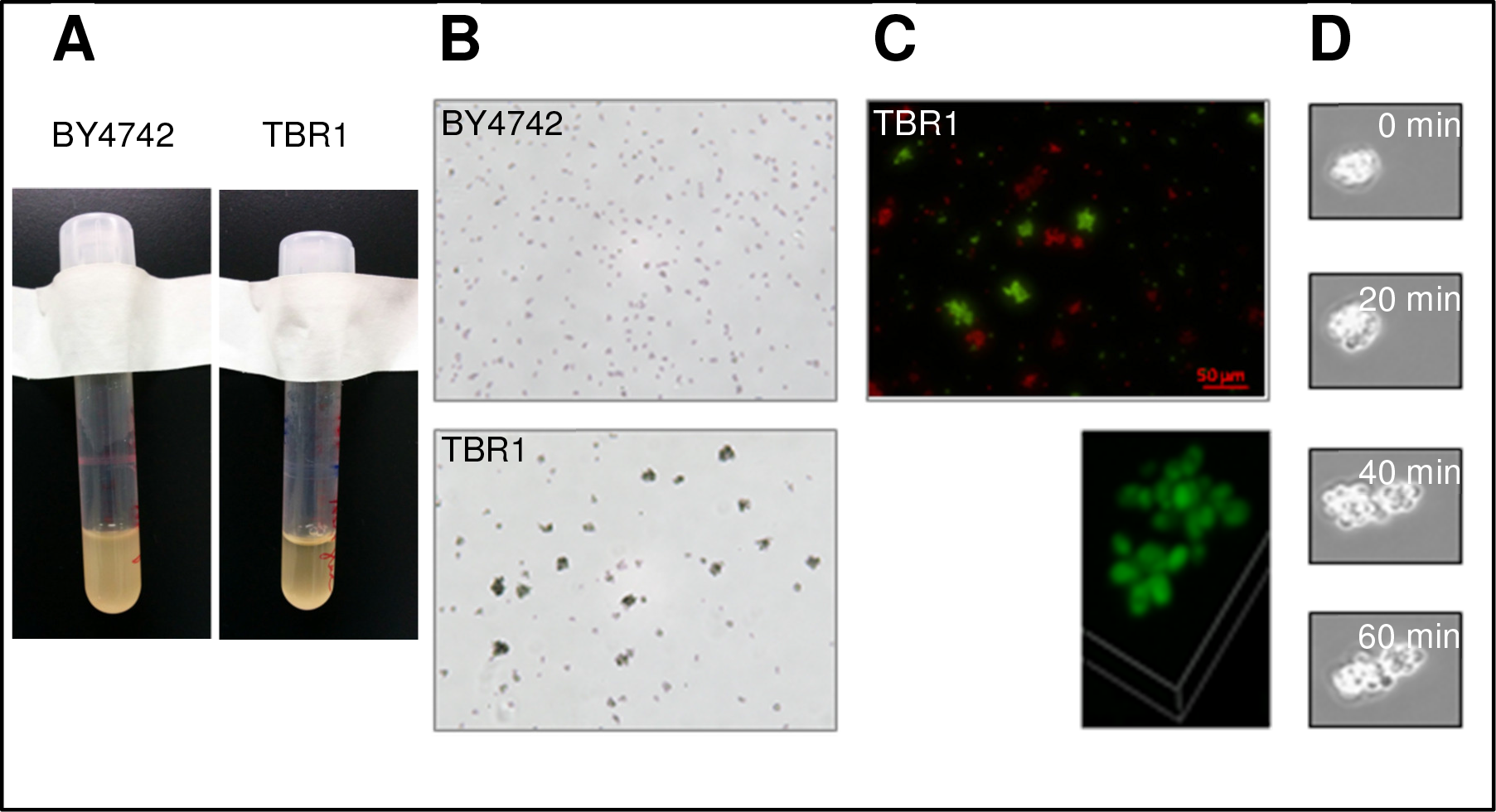
TBR1 cells form clumps by incomplete separation. (A) Settling patterns, 45 minutes after shaking. Yeast cells were kept in YPD + 2% glucose and shaken overnight at 300 rpm at 30°C, moved to a static incubator for 45 minutes, and imaged. (B) Non-clumping laboratory strain BY4742 (top) and clumpy TBR1 strain (bottom) imaged on a Nexcelom Cellometer M10 at 10x magnification. (C) Lack of aggregation in TBR1. Two genetically identical TBR1 strains were tagged with the yEGFP and mCherry reporters expressed from the GAL1 promoter. A 50/50% mixture was resuspended daily in appropriate SD medium for auxotroph selection + galactose. After 7 days, clumps were either purely red or green, indicating lack of aggregation (Nikon TE2000, 20x). Small panel: confocal image, 60x magnification. (D) Clump division: A TBR1 clump divides into two new clumps in static medium, without shaking. Images were taken 20 minutes apart in a Nikon Biostation CT, at 40x magnification.

Besides clump formation by incomplete separation, aggregation (flocculation) balanced by splitting/shedding is another mechanism that could give rise to the multicellular structures seen in Figure 1B. If this were the case then two differently labelled clones should mix within multicellular structures over time as the structures stick together (Smukalla et al. 2008). To test this, we labelled two clones of the clumping strain TBR1 with chromosomally integrated red and green fluorescent reporters, and followed approximately equal numbers of red and green cells over time by fluorescence microscopy. We observed no color-mixing within multicellular structures even after 7 days of co-culture (Figure 1C), indicating that TBR1 cells grow as clumps by incomplete separation, and do not flocculate by aggregation.

Diploid multicellular “snowflake” yeast were found to divide as multicellular units, splitting into two smaller “snowflakes” (Ratcliff et al. 2012). To see if this applied to TBR1 clumps, we tested how clumps multiply without shaking, to avoid the shedding of single-cells due to physical shear. The TBR1 strain existed nearly exclusively in the form of clumps and grew by clump (rather than single-cell) division in these conditions (Figure 1D). This further supported the notion that the TBR1 haploid strain predominantly exists in the form of multicellular clumps that originate from incomplete daughter cell separation and divide as units.

### Multicellular to unicellular transition by laboratory evolution

Next, we asked if TBR1 clumps can undergo a reverse evolutionary transition to unicellularity. Such a transition should result in non-clumping cells that are genetically as similar as possible to their clumping ancestors. These evolved cells would also facilitate testing the benefits and potential costs of clumping during environmental stress: being genetically similar to the ancestors should minimize (although not eliminate) contributions to stress resistance from mechanisms and genes unrelated to clumping.

To induce a reverse transition and obtain two yeast strains that differ in their clumping phenotype but are otherwise genetically similar, we derived a non-clumping, unicellular cell line from the “ancestral” TBR1 strain by laboratory evolution. In 2012, Ratcliff and colleagues selected for multicellular diploid “snowflake” yeast by continuously propagating the yeast that settled most rapidly to the bottom of their cultures (Ratcliff et al. 2012). Here we reversed that selection process by continuously selecting for single-cells or smaller clumps from the tops of our cultures, which remained suspended after the larger clumps settled over 45 minutes (Figure 2A). We called these evolving cell lines “EvoTop”. We also propagated in parallel “EvoControl” cell lines that had the same TBR1 ancestor, but were mixed thoroughly by vortexing before every resuspension. To assess the effect of the counter-gravitational selection for unicellularity, we followed the average size (diameter) of uni- and multicellular structures. A one-way ANOVA revealed significant effects of selection (*F*(2, 6) = 23.91, *p* = 0.001) on TBR1 clump diameters. A Games-Howell post-hoc test indicated that after 4 weeks of daily selection, the EvoTop cell lines formed significantly smaller clumps (*M* = 7.12 μm, *SD* = 0.37) than their ancestors (*M* = 9.88 μm, *SD* = 0.80; *p* = 0.029) or randomly chosen EvoControl lines (*M* = 9.31 μm, *SD* = 0.15; *p* = 0.008) (Figures 2, **S1**), suggesting a reverse evolutionary transition toward unicellularity under the counter-gravitational selection. EvoControl clump size did not differ significantly from ancestral clumps (Games-Howell post-hoc test, *p* = 0.547), but EvoControl clumps tended to be smaller. This could potentially indicate an advantage for smaller clump size even without selection from the top of cultures (Figure 2C).

**Figsure 2.**
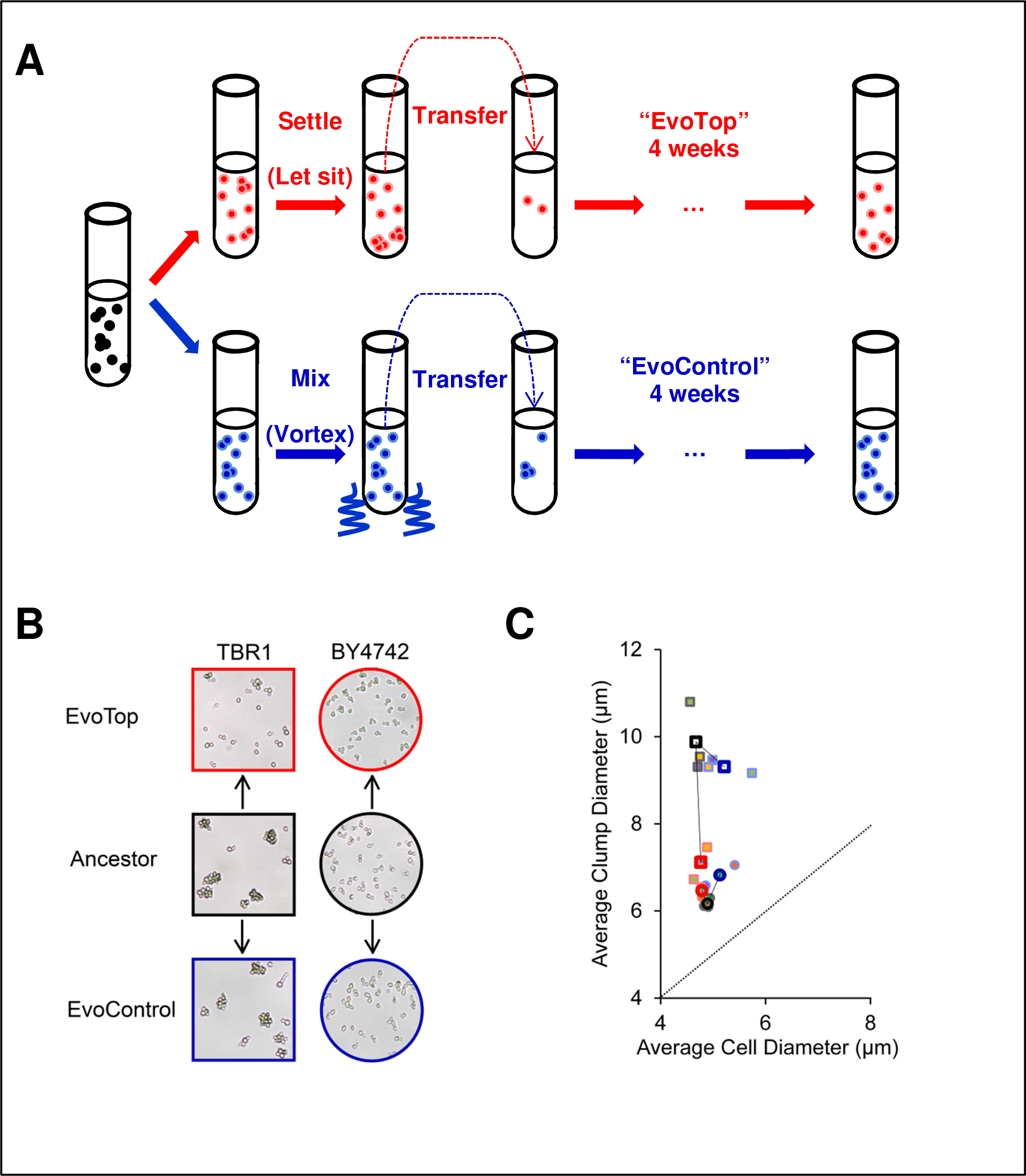
Experimental evolution of unicellularity. (A) Experimental evolution procedure repeated daily over 4 weeks, after ancestral cells were split into “EvoTop” and “EvoControl” cells on the first day. “EvoTop” cells were vortexed and then allowed to settle for 45 minutes before resuspending a small sample from the top of the liquid culture. “EvoControl” cells were vortexed before resuspending the same way. (B) Representative images of TBR1 and BY4742 EvoTop and EvoControl cells compared to their ancestors. The shapes and colors of image frames correspond to the shapes and colors in panel (C). (C) Clump size (diameter) of TBR1 EvoTop (squares outlined in red) decreased compared to the TBR1 ancestor (squares outlined in black) and TBR1 EvoControl (squares outlined in blue) (TBR1 A (purple filled squares), TBR1 B (green filled squares), TBR1 C (orange filled squares)): one-way ANOVA, *F*(2, 6) = 23.91, *p* = 0.001; Games-Howell post-hoc tests, *p* < 0.05. In contrast, BY4742 EvoTop cluster diameters (circles outlined in red) were larger than BY4742 ancestor (circles outlined in black), but this was not significant at a 0.05 alpha level: one-way ANOVA, *F*(2, 6) = 11.53, *p* = 0.009; Games-Howell post-hoc tests, *p* = 0.088. BY4742 EvoControl clusters (circles outlined in blue) were significantly larger than BY4742 ancestral clusters (circles outlined in black): one-way ANOVA, *F*(2, 6) = 11.53, *p* = 0.009; Games-Howell post-hoc test, *p* < 0.05. BY4742 A (purple filled circles), BY4742 B (green filled circles), and BY4742 C (orange filled circles) clump and cell diameter data are also shown. The average cell diameter did not vary significantly with treatment within TBR1 (one-way ANOVA *F*(2, 6) = 3.18, *p* = 0.114) or within BY4742 (one-way ANOVA *F*(2, 6) = 3.40, *p* = 0.103). The dotted line indicates a 1:1 cell diameter to clump diameter ratio, where non-budding unicellular strains would theoretically lie.

There are at least two possible ways for clump size to decrease during evolution: clumps could contain fewer or smaller cells. To distinguish between these possibilities, we compared the cellular diameters of ancestral, EvoControl and EvoTop cell lines. A one-way ANOVA found no significant effects of selection (*F*(2, 6) = 3.18, *p* = 0.114) on the cell diameters of EvoControl lines (*M* = 5.21 μm, *SD* = 0.46), ancestral lines (*M* = 4.67 μm, *SD* = 0.10,) and EvoTop lines (*M* = 4.77 μm, *SD* = 0.13) (Figure 2C). Estimating the average number of cells per clump by dividing the average volume of “spherical” clumps by the average volume of “spherical” cells indicates that the decrease in the TBR1 “EvoTop” clump sizes was driven by an ~2.8 fold decrease in average cell number from 9.5 to 3.3 cells per clump at full packing density (=1), rather than smaller cell size within clumps. This also holds true for TBR1 “EvoControl” clumps, where the average number of cells per EvoControl clump was ~1.7 times smaller than that of the ancestor, at 5.7 cells per clump versus 9.5. Isolates from the TBR1 EvoTop lines exhibited phenotypes approaching that of the BY4742 unicellular laboratory strain (Figures 2B,C; **S1A,B**); therefore, we refer to them as “unicellular” EvoTop TBR1 cells. To test the robustness of these findings, we repeated the evolution experiment for KV38, another clumping strain (Smukalla et al. 2008). Similar to TBR1, the number of cells per clump decreased for EvoTop KV38 lines, confirming that our selection procedure could cause reverse transitions to unicellularity in different yeast cell lines (Figure S1C).

Finally, as a control, we also conducted an identical selection experiment on the standard laboratory strain BY4742. As opposed to TBR1 and KV38, the ancestral cultures of this unicellular strain contained mainly single-cells, intermixed with occasional multiplet structures of two or three cells (doublets or triplets). As expected, the mean “multiplet diameter” of the EvoTop lines did not decrease over time (Figure S1). Surprisingly, however, the presence of small multiplets (mainly doublets and triplets) increased over time in the BY4742 EvoControl and EvoTop. A one-way ANOVA indicated a significant effect of selection (*F*(2, 6) = 11.53, *p* = 0.009) on clump (multiplet) diameters. A Games-Howell post-hoc test revealed that BY4742 EvoControl multiplets (*M* = 6.83 μm, *SD* = 0.23) were significantly larger than BY4742 ancestral multiplets (*M* = 6.17 μm, *SD* = 0.11, *p* = 0.047; Figure 2C), but did not differ significantly from EvoTop multiplets (*p* = 0.189). Like BY4742 EvoControl multiplets, BY4742 EvoTop multiplets (*M* = 6.47 μm, *SD* = 0.13) tended to be larger than BY4742 ancestral multiplets, but this difference was not significant at a 0.05 alpha level (Games-Howell, *p* = 0.088). A one-way ANOVA indicated that the effect of selection (*F*(2, 6) = 3.40, *p* = 0.103) on mean BY4742 cell diameter was not significant among EvoControl cells (*M* = 5.13 μm, *SD* = 0.28), ancestral cells (*M* = 4.89 μm, *SD* = 0.06), and EvoTop cells (*M* = 4.79 μm, *SD* = 0.02) (Figures 2B,C & **S1**), so the increase in BY4742 EvoTop and EvoControl “multiplet size” was not driven by an increase in cell size. This trend may suggest some competitive advantage of such multiplets over single-cells under these conditions, such as covering bud scars, which might cause vulnerability during vortexing (Chaudhari et al. 2012).

In summary, by utilizing the settling rate differences amongst clumps of varying size within a population, we were able to select for unicellular yeast from clump-forming haploid ancestors. Surprisingly, even in the absence of settling-based selection, clump size decreased slightly. In addition, we observed a mild tendency towards multiplet formation for the initially unicellular laboratory strain.

### Genetic bases of unicellularity

To determine the genetic changes underlying the multicellular-to-unicellular transition, we performed whole-genome sequencing on cells from three colonies isolated from each TBR1 EvoTop and ancestral line. We found that each TBR1 EvoTop isolate contained unique mutations in the “Antagonist of Mitotic exit Network” (*AMN1*) gene. The *AMN1* gene (Yvert et al. 2003; Li et al. 2013) is part of the *ACE2* regulon (Di Talia et al. 2009) that mediates the forward transition to multicellularity in yeast (Ratcliff et al. 2015). The TBR1 A EvoTop isolate contained both a loss-of-start missense mutation (Met1Arg) and a stop-gained nonsense mutation (Ser20*), which were present at approximately 72%, and 18% in the population, respectively. We did not observe any differences in the clump-forming abilities over individual experiments that we started from single colonies of the TBR1 A EvoTop isolate, probably because both mutations should effectively knock out the gene, resulting in complete loss of Amn1p function and identical phenotypes. The TBR1 B EvoTop isolate contained a missense mutation in *AMN1* resulting from a Thr405Arg polymorphism. The TBR1 C EvoTop isolate also had a missense mutation in the *AMN1* gene, causing a Lys496Asn amino acid change. In addition to the *AMN1* mutation, the TBR1 B EvoTop isolate also had a Gln1442Leu missense mutation in the adenylate cyclase *CYR1* gene. Information on other mutations identified in the TBR1 EvoTop isolates can be found in Table S1. While the effects of the *AMN1* mutations in TBR1 EvoTop isolate B (Thr405Arg) and C (Lys496Asn) are harder to predict, they likely resulted in decreased or lost function, given the dramatic shift in clump-forming to “unicellular” phenotypes we observed (Figure S2). Taken together, these findings suggest an essential role of the *AMN1* gene in the transition to unicellularity, which is consistent with the known role of Amn1p in regulating a cytokinesis gene network (Wang et al. 2003).

Finally, to investigate the possible role of the *AMN1* sequence in the unicellularity of laboratory strains, we also compared the published sequences of the ancestral TBR1 and the standard laboratory strain BY4742. This also revealed a Asp368Val polymorphism that changes an acidic residue to a hydrophobic one and likely impairs the functionality of the Amn1p protein (Yvert et al. 2003).

### Unicellularity weakens stress resistance, but aids growth in normal conditions

Next, we asked how unicellularity affects stress resistance compared to multicellularity. To compare the stress response of single-celled and clump-forming strains, we isolated three colonies from each TBR1 EvoTop and ancestral line that typified their respective unicellular and clump-forming phenotypes (Supplement, Figure S4). Then we exposed each of these TBR1 isolates to three different stresses: rapid freeze-thaw, 150 μM hydrogen peroxide, and 1% exogenous ethanol, and measured their growth relative to unstressed cells over 1.5 hours. We also similarly compared the relative growth of BY4742 EvoTop and ancestor cells (Figure 3A).

**Figsure 3.**
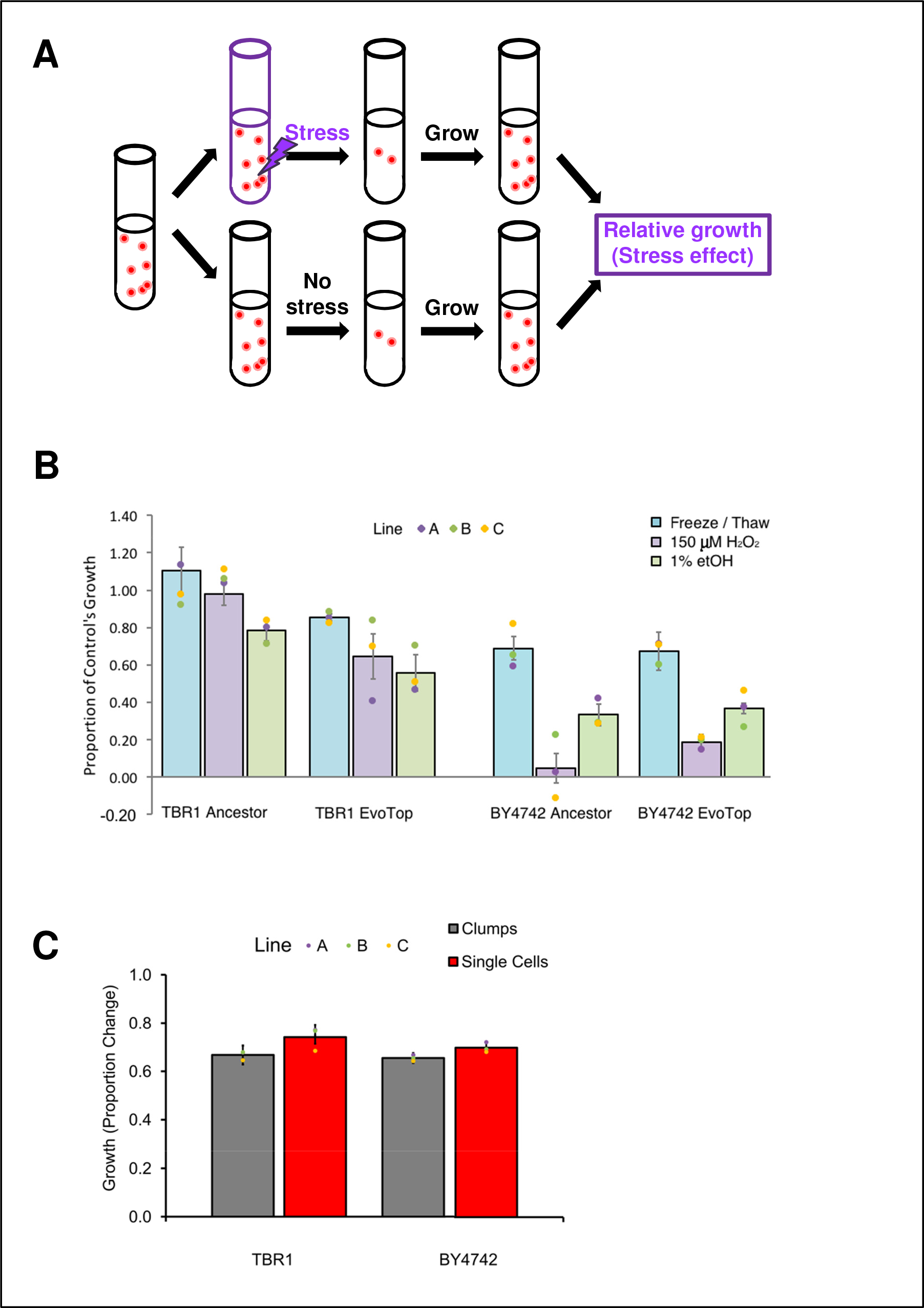
Clumping protects yeast from stress, but hinders growth in normal conditions. (A) Experimental procedure for testing the effects of 3 different forms of stress. Cells were diluted to equal concentrations, exposed to each stress, and then allowed to grow for 1.5 hours. The stress effect (relative growth) was the ratio of growth with stress to growth without stress. (B) Significant differences in the relative growth of stressed cells compared to unstressed controls indicated that single-celled TBR1 EvoTops were less stress resistant than the ancestral clump-forming TBR1 cells (3-way strain (*F*(1, 36) = 25.11, *p* < 0.0001) by stress by line ANOVA, error bars indicate SEM). In contrast, the relative growth of BY4742 ancestral and BY4742 EvoTop strains did not vary significantly from each other after stress treatment (*F*(1, 36) = 1.42, *p* = 0.241). (C) Under stress-free conditions, clump-forming cells had significantly lower growth (*M* = 0.661, *SD* = 0.164) than single-cells (*M* = 0.720, *SD* = 0.129), but growth rates did not vary significantly from each other by strain or line (3-way strain by phenotype (*F*(1, 96) = 4.09, *p* = 0.046) by line ANOVA; error bars indicate SEM).The clump-forming ancestral TBR1 cells had lower growth than EvoTop TBR1 single cells, while ancestral BY4742 unstressed single cells had higher growth than their clumpier EvoTop BY4742 counterparts.

A 3-way strain (TBR1 ancestor, TBR1 EvoTop) by stress treatment (freeze/thaw, 1% ethanol, 150 μM H_2_O_2_) by line (A, B, C) ANOVA revealed significant main effects of strain (*F*(1, 36) = 25.11, *p* < 0.0001) and stress treatment (*F*(2, 36) = 10.91, *p* = 0.0002) on relative growth of stressed cells (Figure 3B). There was not a significant main effect of line on relative growth of stressed cells (*F*(2, 36) = 0.59, *p* = 0.560) or any significant interactions. TBR1 EvoTop (single-celled) lines grew significantly slower (*M* = 0.687, *SD* = 0.220) than the TBR1 ancestral clumping strains (*M* = 0.955, *SD* = 0.237) over 1.5 hours of exposure for all stresses. Therefore, the TBR1-derived single-celled “EvoTop” lines are less resistant to freeze-thaw, hydrogen peroxide, and ethanol stressors than the clumping TBR1 ancestral strains (Figure 3B).

Freeze/thaw stress treatment cells (*M* = 0.979, *SD* = 0.237) had significantly higher relative growth rates than 1% ethanol (*M* = 0.672, *SD* = 0.212; Tukey post-hoc test, *p* = 0.0001) or 150 μM H_2_O_2_ (*M* = 0.812, *SD* = 0.258; Tukey, *p* = 0.041) treated cells, which did not vary significantly from each other (Tukey, *p* = 0.096). We ruled out the potential growth effect of alcohol as a nutrient by growing the cells in media containing 2% glucose, which yeast cells strongly prefer for feeding compared to alcohol (which is the basis of winemaking). Overall, these results implied that clumping provides the benefit of protecting yeast cells from three different forms of environmental stress.

Next, we sought to separately determine if the experimental evolution procedure could somehow lower the stress response of EvoTop lines. To test this, we asked if the single-celled BY4742 ancestral and EvoTop lines differed in their stress resistance. A three-way strain (BY4742 ancestor, BY4742 EvoTop) by stress treatment (freeze/thaw, 1% ethanol, 150 μM H_2_O_2_) by line (A, B, C) ANOVA revealed a significant main effect of stress (*F*(2, 36) = 55.43, *p* < 0.0001), but no main effects of strain (*F*(1, 36) = 1.42, *p* = 0.241) or line (*F*(2, 36) = 0.09, *p* = 0.918) on relative growth of stressed cells (Figure 3B). We found no significant interactions. BY4742 did have significantly higher growth rates after freeze/thaw treatment (*M* = 0.683, *SD* = 0.203) than 1% ethanol (*M* = 0.352, *SD* = 0.106; Tukey, *p* < 0.0001) or 150 μM H_2_O_2_ (*M* = 0.117, *SD* = 0.176; Tukey, *p* < 0.0001) treatment, which were significantly different from each other (Tukey, *p* = 0.0003).

However, we found that BY4742 ancestral and evolved single-cells were similarly vulnerable to freeze/thaw, hydrogen peroxide, and ethanol stress (Figure 3B). Consequently, the unicellular selection procedure, in and of itself, is unlikely to cause increased sensitivity in EvoTop lines.

Finally, to investigate if the single-celled phenotype may also have some benefits over multicellularity, we compared the growth of TBR1 and BY4742 single-cells and clumps in the absence of stress. While the overall growth of TBR1 (*M* = 0.704, *SD* = 0.180) and BY4742 (*M* = 0.677, *SD* = 0.112) did not differ significantly from each other, a 3-way strain (*F*(1,96) = 0.87, *p* = 0.354) by phenotype by line (*F*(4,96) = 0.55, *p* = 0.696) ANOVA revealed a significant main effect of phenotype (*F*(1, 96) = 4.09, *p* = 0.046) on the growth rate of the unstressed cells (Figure 3C). There were no significant strain X phenotype (*F*(1, 96) = 0.25, *p* = 0.621) or phenotype X line (*F*(4, 96) = 0.10, *p* = 0.983) interactions. Single-cells (*M* = 0.720, *SD* = 0.129) divided significantly faster than clump-forming cells (*M* = 0.661, *SD* = 0.164). Accordingly, TBR1 EvoTop single-cells (*M* = 0.741, *SD* = 0.142) grew faster than their clump-forming TBR1 ancestral cells (*M* = 0.668, *SD* = 0.207) in normal conditions. Likewise, unstressed BY4742 ancestral single-cells (*M* = 0.700, *SD* = 0.113) had slightly higher growth than their unstressed clumpier EvoTop counterparts (*M* = 0.655, *SD* = 0.108; Figure 3C). Overall, we concluded that clumping hampered growth, and multicellularity could be costly in the absence of stress. Implicitly, then unicellularity is beneficial without stress.

In summary, the multicellular clump-forming yeast phenotype offers the benefit of increased stress resistance over single-cells. Conversely, we found that clumping might be costly in normal conditions, arguing for the advantage of unicellularity over multicellularity in the absence of stress.

### A simple mathematical model relates clumping to bet-hedging

The experimental findings suggested that clump formation is disadvantageous in the absence of stress, while being beneficial in various forms of stress. To explain these observations and conceptualize the downsides and benefits of yeast multicellularity versus unicellularity, we formulated a simple mathematical model (see the Supplementary Material). The key assumption of this model was that cells closer to the surface of the clump differ phenotypically from the cells hidden inside the clump. Possible sources of such phenotypic differences could be the limited diffusion of nutrients and stressors into the clump interior (Koschwanez et al. 2011), replicative aging (Ratcliff et al. 2015), cell surface area exposed to the growth medium, or other unknown factors.

To model clump-formation, we assumed that cells of radius *r* form spherical clumps of radii *R* (Figure 4A), giving rise to exposed “E” cells on the clump surface and hidden “H” cells in the clump interior. Based on these considerations, the number *n_E_* of exposed “E” cells on each clump’s surface should be roughly proportional to the volume of a shell (dark grey ring in Figure 4A) of width 2*r*. The rest of the cells will be internal or hidden “H” cells buried inside each clump.

**Figsure 4.**
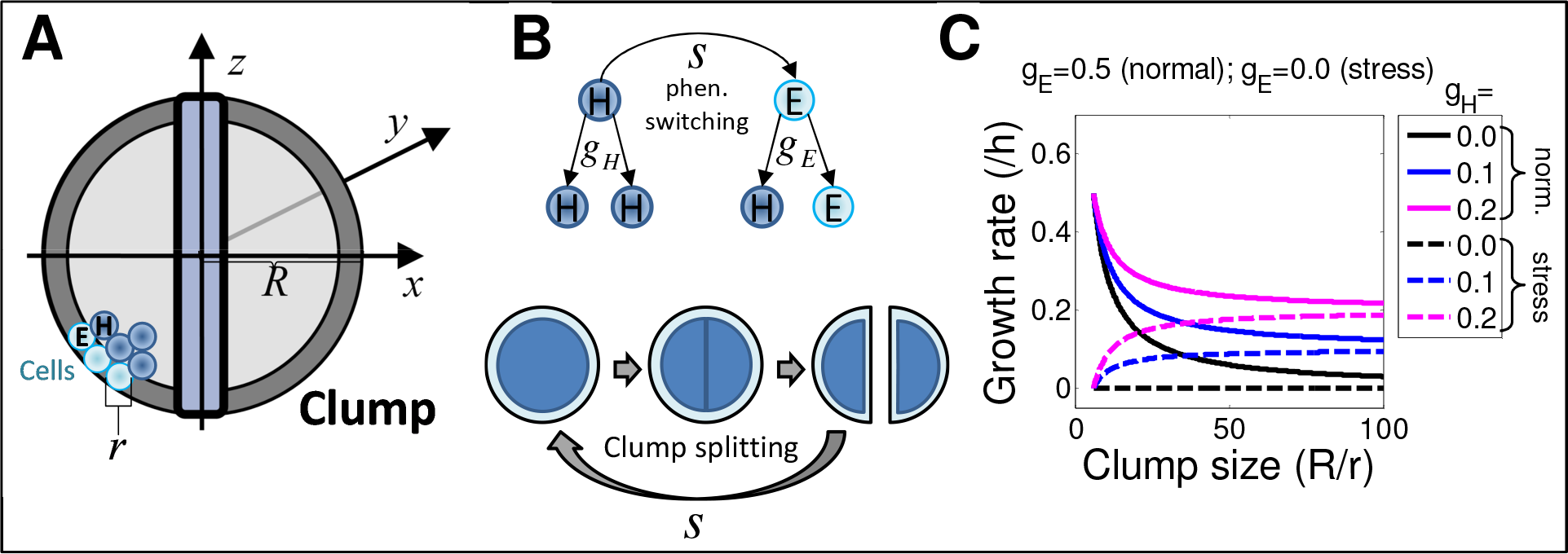
A simple mathematical model of phenotypic heterogeneity in clumping cell populations. (A) The model treats clumps as perfect spheres that have exposed “E” cells on their surface and hidden “H” cells in their interior. (B) Cells grow and switch their phenotype while clumps split through their middle, exposing interior “hidden” cells, effectively causing a phenotypic conversion from “H” to “E” cell type. (C) The graphs with continuous lines represent growth in normal (stress-free) conditions while the ones with dashed lines represent growth in stressful conditions. Calculations show that large clump size is disadvantageous in normal conditions, but beneficial during stress, as observed experimentally.

We postulated that the growth rates of “E” and “H” cells are different and environment-dependent. Specifically, we assumed that “E” cells should grow faster in normal conditions, possibly because they are younger or have direct access to nutrients in the growth medium. On the contrary, the same effects should make “E” cells more vulnerable to stress, so they should grow slower or die more frequently under stress than the hidden “H” cells. To estimate how cells may transition between “E” and “H” states as they grow at environment-dependent rates *g_E_* and *g_H_*, respectively, while clumps split at rate *s*, we considered that when a mother clump divides, “H” cells become exposed on both sides of an equatorial division surface (Figure 4B). Assuming that each exposed cell gives rise to a hidden cell during clump growth, while hidden cells give rise to two hidden cells, the following equations (see the Supplementary Material) capture the division of clumps and single-cells, as well as phenotypic fluctuations as hidden cells become exposed:

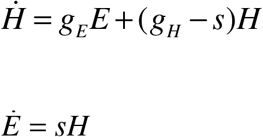

A short calculation (see the Supplementary Material) leads to the conclusion that in the absence of stress, the growth rate will decrease with the clump size *R* as shown in Figure 4C. Conversely, the growth rate will increase with the clump size when stress is present. These predictions explain the experimental observations in Figure 3B,C. Therefore, this mathematical model intuitively illustrates how clump formation might create phenotypic heterogeneity and phenotype switching in isogenic cell populations, giving rise to “E” and “H” cell types. This phenotypic heterogeneity is beneficial in stress, but disadvantageous in normal conditions. Since “E” cells grow faster than “H” cells in normal conditions, while the reverse is true in the presence of stress, the phenotypic effects of clumping amount to regulated (as opposed to random) phenotype switching (Aldridge et al. 2012), causing bet-hedging diversification that may ensure survival during unexpected periods of stress (Acar et al. 2008; Levy et al. 2012).

## 4. DISCUSSION

While the transition from unicellularity to multicellularity has been studied extensively, the reverse transition (from multicellularity to unicellularity) has been less investigated. We obtained a reverse transition by EvoTop selection, as a new exception from Dollo’s Law (Dollo 1893). We found that clumping protects from stresses but is costly in normal conditions, suggesting that clumping should also be lost without selection, in the absence of stress. Indeed, TBR1 clump size decreased not only in EvoTop lines, but also slightly in EvoControl lines during experimental evolution. This may imply a trade-off between growing fast and dying in stress (unicellular phenotypes) versus growing slower, but resisting stress (multicellular phenotypes). A similar trade-off was recently observed in the evolution of engineered unicellular yeast cells under stress (Gonzalez et al. 2015). Such trade-offs emerge from the pressure to satisfy two conflicting tasks, resolved by Pareto optimality in biological evolution (Shoval et al. 2012). Overall, these findings suggest that environmental stress could play a major role in the maintenance, or possibly even the emergence of multicellularity in other species, such as social amoebae (Gregor et al. 2010). These findings corroborate the protective role that clumping or other forms of multicellularity provide against predation (Brunke et al. 2014; Pentz et al. 2015) and environmental stress (Smukalla et al. 2008).

We found that that haploid yeast clumps are more resistant than single-cells against both physical (freeze/thaw) and chemical (ethanol and hydrogen peroxide) stressors. Interestingly, protection from freeze/thaw appears to be a benefit that the flocculating multicellular form does not possess (Smukalla et al. 2008). Strikingly, not only did the TBR1 ancestral cells resist freeze/thaw treatment, but they grew ~110% better than their corresponding untreated controls. An explanation is that freeze/thaw cycles impose mechanical stress (Harju et al. 2004), which could cause clump splitting, improving access to nutrients. Interestingly, the unicellular BY4742 strain also started forming more multiplets during evolution, which may have been selected for because they mitigate bud scars’ vulnerability to mechanical stress from vortexing (Chaudhari et al. 2012). The higher stress tolerance of clumps may be due to either physical shielding of interior cells from external stressors, replicative aging, physiological changes, or a combination of such factors (Brachmann et al. 1998; Smukalla et al. 2008).

Our findings and mathematical model jointly suggest that clumping may create phenotype switching and deterministic heterogeneity, which plays a role in the drug resistance of microbial pathogens (Aldridge et al. 2012). Indeed, some *S. cerevisiae* strains emerging as opportunistic pathogens (Wei et al. 2007) contain the same *AMN1* allele as the clump-forming TBR1 ancestor. Moreover, long-term evolution of *Candida glabrata* with macrophages causes the evolution of a filamentous multicellular form due to incomplete daughter cell separation (Brunke et al. 2014). Further work is needed to examine what role clump formation might play in yeast infections, but one can envision clumps forming stress resistant propagules. Future studies of clumping and other forms of multicellularity in infectious yeasts and other microbes should lead to improved antibiotic efficiency, addressing the emerging global threat of drug resistant infections.

## ACKNOWLEDGEMENTS

We would like to thank the members of the Balázsi laboratory and Dr. Gerda Saxer, Dr. Chris Kuzdzal Fick, and Dr. Michael Lorenz for discussions. We also thank Dr. Kevin Verstrepen, and Dr. Todd B. Reynolds for strains.

## SUPPORTING MATHEMATICAL MODEL

### Cellular dynamics in clumping populations

To develop a simple model that could capture the phenotypic effects of clump formation, we considered cells of radius *r* that form multicellular spheres (clumps) of radius ***R***. We assume that cells on the surface of these spheres are phenotypically different from the interior: they divide faster in normal conditions, while the opposite is true in environmental stress conditions. Our first goal will be to roughly estimate the fraction of cells on the surface versus inside the spheres, assuming a filling factor =1 and that all cells are identical in size.

To estimate the number of cells per clump, we start from the volume of each sphere (clump):

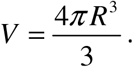

Then we can estimate the total number of cells (of volume *v*) in such a spherical clump given by:

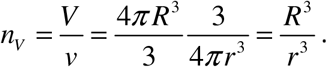

To estimate the number of exposed “E” cells, we consider that the surface of the sphere is:

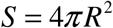

The number of “E” cells near the surface of the sphere can be estimated based on the volume of a shell whose width is the diameter 2*r* of a single-cell:

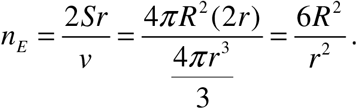

Consequently, the approximate number of hidden “H” cells is equal to:

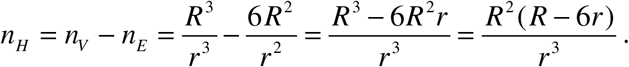

From here, the fractional ratios of cells on the surface versus the interior are, respectively:

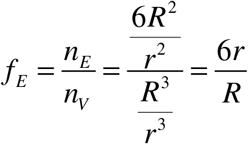

and

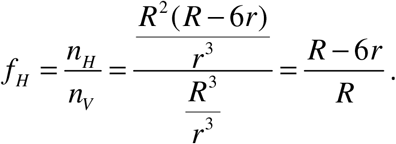

The ratio of cells on the surface versus in the interior will be given by:

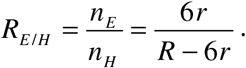

Next, we analyze how cells grow and switch phenotype by transitioning between clump interiors and exteriors in the absence of stress. Over each clump life cycle (between two clump splittings) we consider two phases of population dynamics: (i) phenotype switching that predominantly occurs during clump splitting and (ii) growth by cell divisions that predominantly happen before clump splitting. First, we analyze phenotype switching during clump division.

When the clump divides, it exposes its equatorial cross-section, which has a surface area

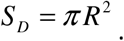

The total number of cells on both sides of the division plane can be estimated as:

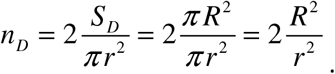

The chance for a hidden cell to end up on the division plane (and thus become exposed) is:

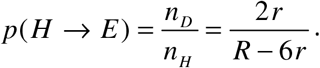

Consequently, the chance for a hidden cell to remain interior during clump division is:

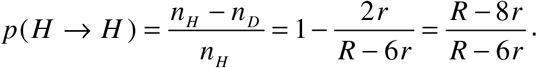

In addition, the chance for an exterior cell to internalize during division is 0. This further implies that the chance for an exposed cell to stay exposed at clump division (splitting) is 1.

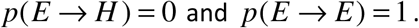

Overall, phenotype switching over the clump’s life cycle (lasting for time Δ*T* between two clump splitting events) will be described by a difference equation involving a transition matrix:

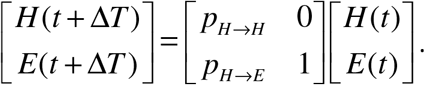

The right-hand side can be split into two parts:

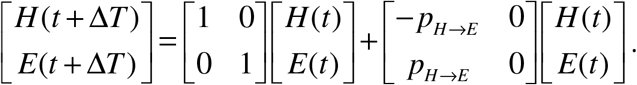

With a little algebra we obtain the equation for phenotypic switching:

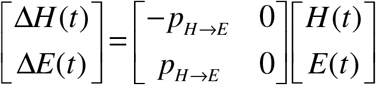

In addition to phenotypic switching, hidden and exposed cells can divide between two clump splitting events. We assume that E cells give rise to an E cell and an H cell while growing at rate *g_E_*. Hidden H cells only give rise to two H cells as they divide while growing at rate *g_H_*. Overall, the growth and switching dynamics for clumping cell populations can be roughly approximated over the time period Δ*T* between two splitting events with the following set of equations:

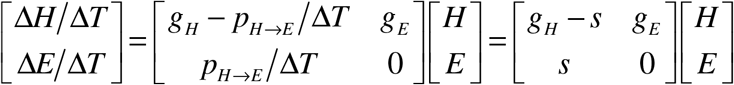

In the above equation we denoted the phenotypic switching rate as:

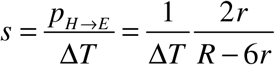

Overall, population dynamics is governed by a system of two ordinary differential equations:

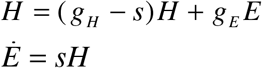

For a sanity check, we add the two equations to obtain: *Ḣ* + *Ė* = *g*_*E*_*E* + *g*_*H*_*H* = *g*(*H* + *E*).

In the long-term limit, switching and growth balance each other out to give: *H* = *k*_*H*_*e^gt^* and *E* = *k*_*E*_*e^gt^*.

From the second differential equation, we obtain: *Ė* = *gk*_*E*_*e^gt^* = *sH* = *sk*_*H*_ *e^gt^*, which implies: 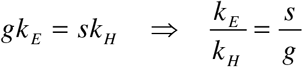.

Also, we should have: 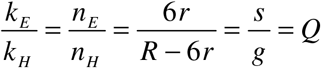.

From the first differential equation, we obtain:

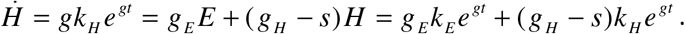

After we rearrange and simplify, we get: 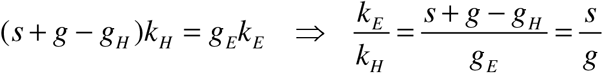.

Using the expression for *Q*, we obtain after some basic algebra:

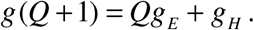

From here, we finally obtain the overall growth rate:

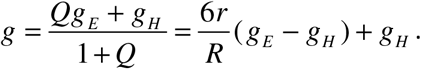

We plot this overall growth rate *g* as a function of *R* for various pairs of *g_E_*, *g_H_* values chosen to represent normal and stressful conditions in Figure 4 of the main text.

## SUPPORTING METHODS AND ANALYSIS

### Fluorescent labelling of TBR1

The integration of GFP and mCherry reporters into TBR1 chromosomes was described previously (Chen et al. 2014). Briefly, *Escherichia coli* strains with either the pDN-G1Gh (GFP) or the pDN-G1Ch (mCherry) plasmids (both of which harbour the ampicillin resistance selection marker), were incubated in LB media with Ampicillin (1:1000) at 37 C for 6-8 hours. The plasmids were then extracted by midi prep (QIAGEN), linearized, and purified. They were then transformed into the native GAL1 locus of the TBR1strain, using the histidine auxotrophic marker. Transformation was performed using a modified lithium acetate procedure as described before (Chen et al. 2014). Synthetic drop-out (SD-his-tryp) plates were then used for selection (all reagents from Sigma, Inc.). Once established, TBR1-GFP and TBR1-mCherry strains were incubated at 30°C, shaking at 300 rpm (LabNet 311DS shaking incubator). The TBR1-GFP and TBR1-mCherry strains were imaged and cells were counted under Nikon fluorescence microscopy and FACScan respectively.

### TBR1 sequencing analysis

The DNA from the ancestor and each of the three TBR1 EvoTop isolates was extracted from cultures started from individual colonies that were anticipated to be clonal. Clonal mutations are expected to be present at a minimum of 80% – 90% in clonal population samples analyzed as if they were polymorphic (Saxer et al. 2014). Since the TBR1 A EvoTop isolate appeared to be founded by two clones, with the lower frequency clone present at about 18%, we reasoned that this clone could have other clonal mutations present at as low as 14% (= 0.8*18%) (Table S1).

We used high throughput whole genome sequencing to identify the mutations underlying the change from clumping to “unicellular” phenotypes in our TBR1 EvoTop lines and isolates. We followed QIAGEN’s “Purification of total DNA from yeast using the DNeasy Blood & Tissue Kit” protocol to obtain high quality genomic DNA from the TBR1 ancestor, EvoControl, and EvoTop lines and isolates. Columbia Genome Center used these samples to run whole genome sequencing on an Illumina HiSeq 2000 v3 instrument, and they aligned the obtained reads in fastq files to the ∑1278b reference genome (Dowell et al. 2010) using BWA-mem.

We prepared the provided aligned bam files for analysis by adapting the Genome Analysis Toolkit’s (GATK 3.3-0) best practices procedures (Mark Duplicates with REMOVE_DUPLICATES=true (Picard-tools 3.2.53(1)), indel realignment (GATK), quality recalibration (GATK)) and added alignment qualities with lofreq_star-2.1.2). Because the EvoControl and EvoTop populations were heterogeneous, we looked for low frequency mutations in the genomic DNA of all 10 samples (TBR1 ancestor, TBR1 A, B, & C EvoControl, EvoTop, and EvoTop isolates) with lofreq_star-2.1.2 (Wilm et al. 2012). In addition, we used the breseq-0.26.1 pipeline (Deatherage and Barrick 2014) with bowtie2-2.2.6 and the −j2, −p, −c options to align fastq files to the ∑1278b reference genome (Dowell et al. 2010) in order to independently identify variants in each of the 10 lines. To identify mutations in the EvoTop isolates that were not called in the ancestor we used the vcf-isec command of VCFtools 0.1.12b with option −c for LoFreq* files and gdtools SUBTRACT for breseq files. We used vcf-isec −f −n +2 to identify which of these mutations were called by both LoFreq* and breseq and then ran LoFreq*’s uniq command to identify which of these mutations were really unique from the ancestor. To help determine the potential impact of these mutations, we used gdtools ANNOTATE and the Sigma1278b_ACVY01000000.gff file deposited at yeastgenome.org by Dowell and colleagues (Dowell et al. 2010). We analyzed all 10 lines as polymorphic populations so that we could directly compare the results. Mutations in the TBR1 EvoTop isolates that were unique from the ancestor and were called by both LoFreq* and breseq with a minimum frequency of 14% (see above) are listed in Table S1.

### Assays of stress resistance

#### Testing for stress resistance: Freeze/Thaw

For the freeze/thaw treatment, we rapidly froze cells in a dry ice and ethanol bath for 5 minutes and immediately thawed them in room temperature water. During this time, the remaining 1.5 ml cell samples stayed on the bench top as controls. We prepared 3X dilutions of each culture by adding 1 ml of cells to 2 ml SD in 5 ml polystyrene round-bottom yeast culture tubes and transferred 1 ml of each sample into new yeast culture tubes. We placed the 2 ml samples back into a 30°C, 300 rpm Labnet 311DS shaking incubator and prepared Nexcelom automated cell counter slides from each ancestral and EvoTop cell line. In order to break up clumps and obtain accurate cell counts, we used a Qsonica Q55 sonicator to administer 20 pulses of 30% amplitude sonication to each of the 1 ml samples, which were placed on ice between each set of 10 pulses. We subsequently acquired clump diameter data before and after sonication, along with cell concentration data with a Nexcelom Cellometer Vision automated cell counter using the same parameters as the Cellometer Auto (with the additional background adjustment parameter for cell measurements set at 1.0). After 1.5 hours of growth in the incubator, we prepared new 1 ml samples of each culture and followed the previously described protocol to obtain clump diameter and cell concentration data. We then calculated the proportion change in cell concentration for the EvoTop single-cells and ancestral clumps under freeze/thaw and control conditions as follows: (final concentration - initial concentration)/ initial concentration. To obtain the relative growth of stressed cells compared to their controls, we divided the proportion change of each freeze/thaw sample by the proportion change in its control counterpart ((proportion change freeze/thaw) / (proportion change control)). We followed the same protocol to examine the effects of freeze/thaw stress on our BY4742 A, B, and C lines, but determined that sonication was not needed to obtain countable cells.

#### Testing for stress resistance: 150 μM H_2_O_2_ and 1% exogenous ethanol

In order to determine if clumps of yeast are more resistant than single-cells to the chemical stressors hydrogen peroxide and ethanol, we followed a similar protocol to the one described for freeze/thaw. We diluted exponentially growing cells started from individual colonies of TBR1 A, B, and C EvoTop and ancestral isolates to 3.5*10^6^ cells/ml in SD. We made 2X dilutions of the cells for control and 150 μM H_2_O_2_ or 1% ethanol treatments by adding 1 ml of cells to prepared 5 ml yeast culture tubes containing 1 ml SD for controls and either 3.41 μl of 0.3% H_2_O_2_ or 2% ethanol in 1 ml SD for stress treatments. To obtain samples for initial counts, we transferred 1 ml of each culture into new yeast culture tubes. Then we placed the remaining samples back into the 300 rpm shaking 30°C LabNet 311DS incubator. We allowed the single-cell and clump lines to grow for 1.5 hours in a 30°C, 300 rpm LabNet 311DS shaking incubator. To calculate the proportion change in concentration of EvoTop single-cells and ancestral clumps, we used sonicated cell counts obtained from a Nexcelom Cellometer Vision automated cell counter in the manner previously described for the freeze/thaw experiment. We also calculated and analyzed the relative growth of stressed cells compared to their controls for H_2_O_2_ or ethanol samples as previously described for the freeze/thaw experiment. We followed the same protocols with our BY4742 A, B, and C EvoTop and ancestral lines, but did not sonicate the cells for counting.

### Selection for the unicellular yeast form of KV38

To test if the observed effects were strain-dependent, we also applied identical counter-gravitational selection to evolve another clumping strain, KV38 a haploid segregant of EM93 (Smukalla et al. 2008). Like TBR1, EvoTop lines of KV38 decreased in clump size (Figure S1 A & C). When we started the selection experiment, the KV38 ancestor did not appear to flocculate, so EDTA was not added to the sample we obtained live ancestral clump and cell size data from. However, KV38’s tendency to flocculate fluctuated throughout the selection experiment, so we added 5 mM EDTA to break up flocs before measuring clump and cell size.

A one-way ANOVA revealed a significant effect of selection (*F*(2, 6) = 28.61, *p* = 0.0009) on clump size. A Games-Howell post-hoc test indicated mean EvoTop clump diameter (*M* = 6.26 μm, *SD* = 0.21) was significantly smaller than that of the ancestral (*M* = 7.77 μm, *SD* = 0.25) and EvoControl lines (*M* = 7.67 μm, *SD* = 0.35) (*p* < 0.05), which did not vary significantly from each other (Figure S2). A one-way ANOVA did not show a significant effect of selection (*F*(2, 6) = 3.93, *p* = 0.0813) on KV38 cell size, indicating that the diameters of EvoControl cells (*M* = 5.62 μm, *SD* = 0.62), ancestral cells (*M* = 4.89 μm, *SD* = 0.05), and EvoTop cells (*M* = 4.93 μm, *SD* = 0.037) were not significantly different from each other. Assuming a packing density of 1, the average ancestral KV38 clumps contained nearly twice as many (~4.01) cells per clump as the KV38 EvoTop (~2.05) strain and 1.58 times as many cells per clump as the EvoControl (~2.54) strain. Therefore, these results indicate that not only KV38 EvoTop, but also KV38 EvoControl clumps were comprised of fewer cells than KV38 ancestral clumps. Along with the TBR1 results, this suggests that having fewer cells per clump might be beneficial under the growth conditions of our experiment, with or even without settling-based selection.

### Histograms of clump diameters

Histograms showing the distribution of clump diameters in SD medium of the TBR1 (isolates) and BY4742 ancestral and EvoTop lines indicate that the distribution of BY4742 clump diameters stayed consistent between the ancestral and EvoTop lines. However, the distribution of TBR1 EvoTop isolate clump diameters shifted away from the TBR1 ancestral distribution and closer to that of BY4742 (Figure S1B). Each TBR1 isolate shared a similar distribution of clumps (Figure S2). The clump distributions shown in Figure S1B are from the combined data points of all lines (A, B, and C) within a strain (TBR1 ancestor, TBR1 EvoTop, BY4742 ancestor, and BY4742 EvoTop). The ancestral TBR1 clump distribution in Figure S2 is an average of the TBR1 ancestral isolate A, B, and C clump diameter distributions.

### Growth of stress-exposed EvoTop isolates

The TBR1 B EvoTop isolate (*M* = 0.808, *SD* = 0.147) had significantly higher relative growth after exposure to stress than the TBR1 A EvoTop isolate (*M* = 0.575, *SD* = 0.272), but its growth did not vary significantly from the TBR1 C EvoTop isolate (Figure S3). The TBR1 C EvoTop isolate (*M* = 0.677, *SD* = 0.176) tended to have lower stressed growth than the TBR1 B EvoTop isolate, but it did not vary significantly from that of the TBR1 A or B EvoTop isolates (Figure S3).

### TBR1 Isolates

For each TBR1 ancestor and EvoTop strain and line (A, B, C) we took 6 colonies and used each to inoculate 1 mL SD tubes for overnight growth in a shaking 30°C 300 rpm incubator. We froze isolates (700 μL SD with 300 μL 80% glycerol) and prepared dilutions in SD for exponential growth overnight. The following day, we used the Nexcelom Cellometer Vison CBA “Clumpy Yeast” parameters to measure clump diameters, distributions, and concentrations of isolates (10X dilutions, except for TBR1 C ancestor 6, which was 1X). Isolates with extremely low concentrations (< 1*10^6^ clumps/mL) were not considered for further use. We took average clump diameter and clump size distribution into account to determine which ancestral and EvoTop isolates to use in our stress tests. We chose EvoTop isolates with small clumps that lacked larger clumps in their distribution data. For ancestral isolates, we chose ones with large clumps and distributions that lacked very small clumps. Isolates shown in red (Figure S4) were used for stress tests. TBR1 C ancestor 2 (green) was used in the first Freeze/Thaw experiment, but TBR1 C ancestor 5 was used in all other stress test replicates.

### TBR1 EvoTop isolates

The EvoTop isolates from TBR1 B and C were clonal, but our sequencing results indicated that the TBR1 A EvoTop isolate contained 2 clones. The *FLO11* gene of TBR1 encodes a surface flocculin that is required for biofilm cell-surface adhesion and involved in cell-cell adhesion (Lo and Dranginis 1998; Reynolds and Fink 2001). The TBR1 A EvoTop colony we isolated appears to have been founded by cells from 2 genetically distinct lines that may have adhered to each other. For that colony, there are two mutations in the *AMN1* gene that are within the 101 bp Illumina HiSeq reads. Each individual read that overlaps both locations contains either one mutation or the other.

### Stress effects in flocs and clumps

Both physical shielding and physiological changes appear to play a role in the increased stress tolerance of flocs compared to single-cells (Smukalla et al. 2008). Along with being blocked from external stressors by exterior floc cells, internal floc cells have limited access to nutrients and oxygen. This leads to decreased growth as indicated by the downregulation of mitotic genes. There is a corresponding upregulation of genes associated with stress response (Smukalla et al. 2008). Smukalla and colleagues (Smukalla et al. 2008) were able to test the effects of different stressors on flocculating and non-flocculating cells by breaking up flocs with EDTA after exposure to stress and measuring colony forming units. Our clumps formed from incomplete daughter cell separation do not lend themselves well to the colony forming unit assay, which led us to examine the effects of stress by calculating the relative growth of stressed cells compared to unstressed controls. Clumps are not easily broken up, and the mechanical method of sonication we used could lyse and kill the cells. While these cells are still countable in our initial and final time points, they would not show up in a colony forming unit assay. Similarly, very small countable clumps existed in the TBR1 samples after sonication, but these multiple cells would have only formed a single colony. We sonicated samples of the TBR1 cultures that we took to count, but the cells we measured for growth were not exposed to sonication until we were ready to obtain single-cell counts after they had undergone 1.5 hours of growth.

### Mutations in EvoTop Isolates

In addition to the *AMN1* mutations we identified, our TBR1 EvoTop isolate sequencing results revealed a few mutations that could potentially affect stress response (Table S1). The most notable of these is the *CYR1* mutation found in the TBR1 B EvoTop isolate. *CYR1* is an essential gene that encodes adenylate cyclase, and null mutants are inviable (Giaever et al. 2002). The TBR1 B EvoTop isolate was not only viable, but its relative growth after stress exposure was significantly higher than that of the TBR1 A EvoTop isolate, and while not varying significantly, it showed a higher relative growth trend than the TBR1 B EvoTop isolate (Figure S3). This suggests that the *CYR1* mutation may not have had a negative effect on the TBR1 B isolate’s stress tolerance. The TBR1 B EvoTop isolate also had a mutation in *OSW1*, and one intergenic to *YKE4* and *TIM44*. The TBR1 A EvoTop isolate contained mutations in *BAP2*, *DOT6*, and *VPS13*. While none of these mutations are in specific stress response genes, they may ultimately affect stress tolerance in these strains. Interestingly, out of these 5 mutations, only the *DOT6* mutation in the TBR1 A EvoTop isolate was also called in the EvoTop pool it was isolated from. The other mutations were either present at too low a frequency to be picked up on during sequencing, or are truly novel to the isolates. The former is the likely case for the *OSW1* mutation in the TBR1 B EvoTop isolate, which can be found at low frequency (~3%) in the aligned reads from the TBR1 B EvoTop population. There is no evidence of the remaining three mutations in the files from their corresponding pools, but this alone is not enough evidence to determine if the mutations are truly unique to the isolates or were simply at too low a frequency in the pools to show up under our given coverage. Aside from the *AMN1* mutation, we did not identify any other high frequency mutations in the TBR1 C EvoTop isolate. Notably, the relative stressed growth of the TBR1 C EvoTop isolate did not vary significantly from that of the TBR1 A or B EvoTop isolates (Figure S3), and the overall relative stressed growth of the TBR1 EvoTop isolates was significantly higher than that of the BY4742 ancestral and EvoTop lines (Figure 3). This suggests that stress tolerance of the TBR1 EvoTop isolates has not been compromised by random mutations.

**Figsure S1.**
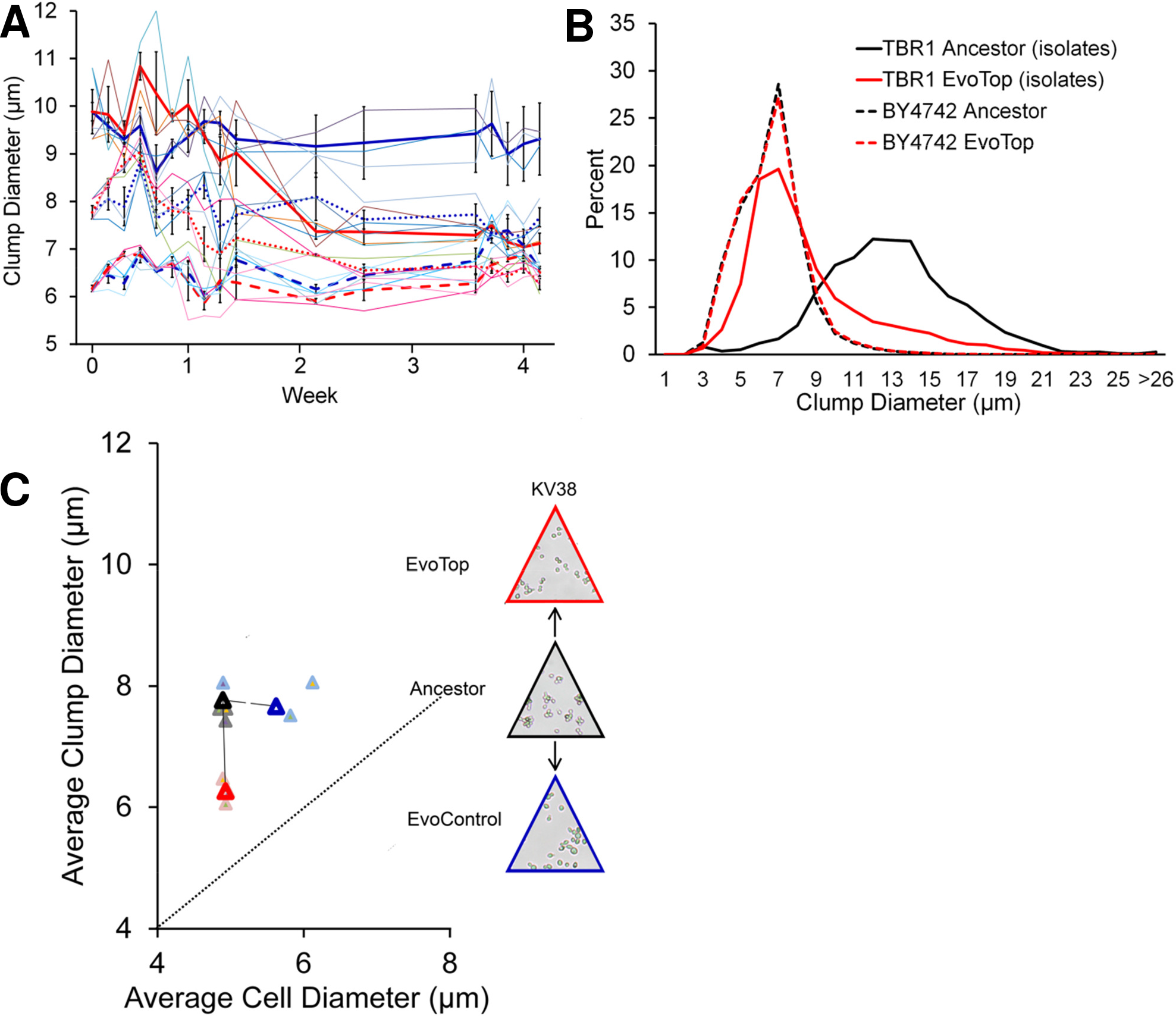
**(A)** TBR1 (solid bold lines) and KV38 (dotted lines) EvoTop clumps (red lines, week 4) were significantly smaller than their corresponding ancestral (week 0) and EvoControl clumps (blue lines, week 4), while the average BY4742 (dashed lines), EvoControl, and EvoTop clump diameters (week 4) tended to be larger than their ancestor’s (week 0). Replicates are shown in thin blue and red lines. Error bars indicate SEM, N=3. **(B)** In SD medium, the clump diameter distribution of TBR1 EvoTop isolates (solid red line) approached that of the unicellular laboratory strain BY4742 (dashed red line). **(C)** KV38’s mean EvoTop (triangle outlined in red) clump diameter (*M* = 6.26 μm, *SD* = 0.21) was significantly smaller than that of the ancestral (triangle outlined in black; (*M* = 7.77 μm, *SD* = 0.25) and EvoControl lines (triangle outlined in blue; *M* = 7.67 μm, *SD* = 0.35) (one-way ANOVA (*F*(2, 6) = 28.61, *p* = 0.001) followed by Games-Howell post-hoc test, *p* < 0.05, error bars indicate SEM), which did not vary significantly from each other (*p* > 0.05). The diameters of EvoControl cells (*M* = 5.62 μm, *SD* = 0.62), ancestral cells (*M* = 4.89 μm, *SD* = 0.05), and EvoTop cells (*M* = 4.93 μm, *SD* = 0.037) did not vary significantly from each other (one-way ANOVA (*F*(2, 6) = 3.93, *p* = 0.081). KV38 A (purple filled circles), KV38 B (green filled circles), and KV38 C (orange filled circles) clump and cell diameter data are also shown. The dotted line indicates a 1:1 cell diameter to clump diameter ratio, where non-budding unicellular strains would theoretically lie.

**Figsure S2.**
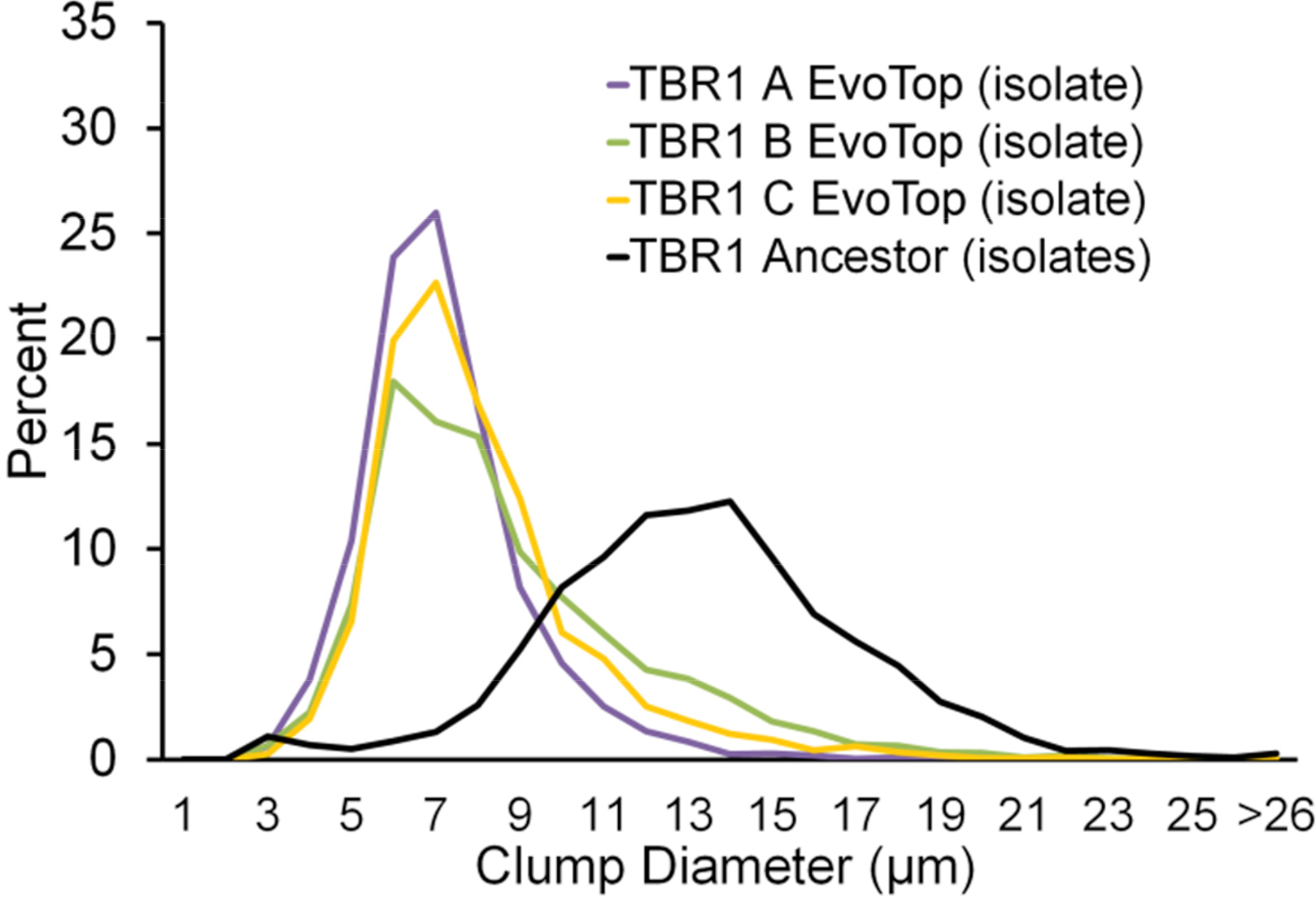
In SD medium, the TBR1 B EvoTop isolate (green line) and the TBR1 C EvoTop isolate (orange line) have clump diameter distributions that are similar to that of the TBR1 A EvoTop isolate (purple line).

**Figsure S3.**
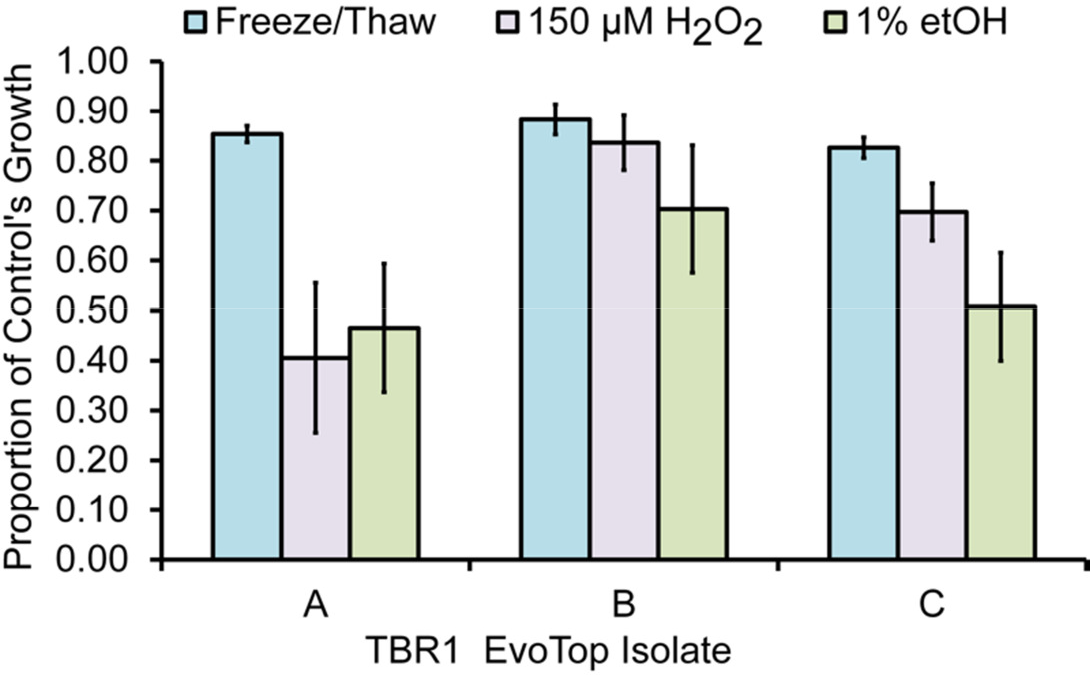
The TBR1 B EvoTop isolate had significantly higher relative growth after stress exposure than the TBR1 A EvoTop isolate (2-way line (*F*(2, 18) = 4.90, *p* = 0.020) by stress treatment (*F*(2, 18) = 8.31, *p* = 0.003) ANOVA followed by a Tukey post-hoc test, *p* < 0.05, error bars indicate SEM). The relative growth of stressed TBR1 C EvoTop isolate cells tended to be lower than that of the TBR1 B EvoTop isolate, but did not vary significantly from the TBR1 B EvoTop isolate or TBR1 A EvoTop isolate (Tukey post-hoc test, *p* > 0.05). Within the EvoTop isolates, freeze/thaw treated cells (blue bars; *M* = 0.854, *SD* = 0.043) had higher relative growth than 150 μM H_2_O_2_ (purple bars; *M* = 0.646, *SD* = 0.241; Tukey, *p* < 0.05) and 1% etOH (green bars; *M* = 0.559, *SD* = 0.213; Tukey, *p* < 0.05) treated cells, which did not vary significantly from each other (Tukey post-hoc test, *p* > 0.05; Statview 5.0).

**Figsure S4.**
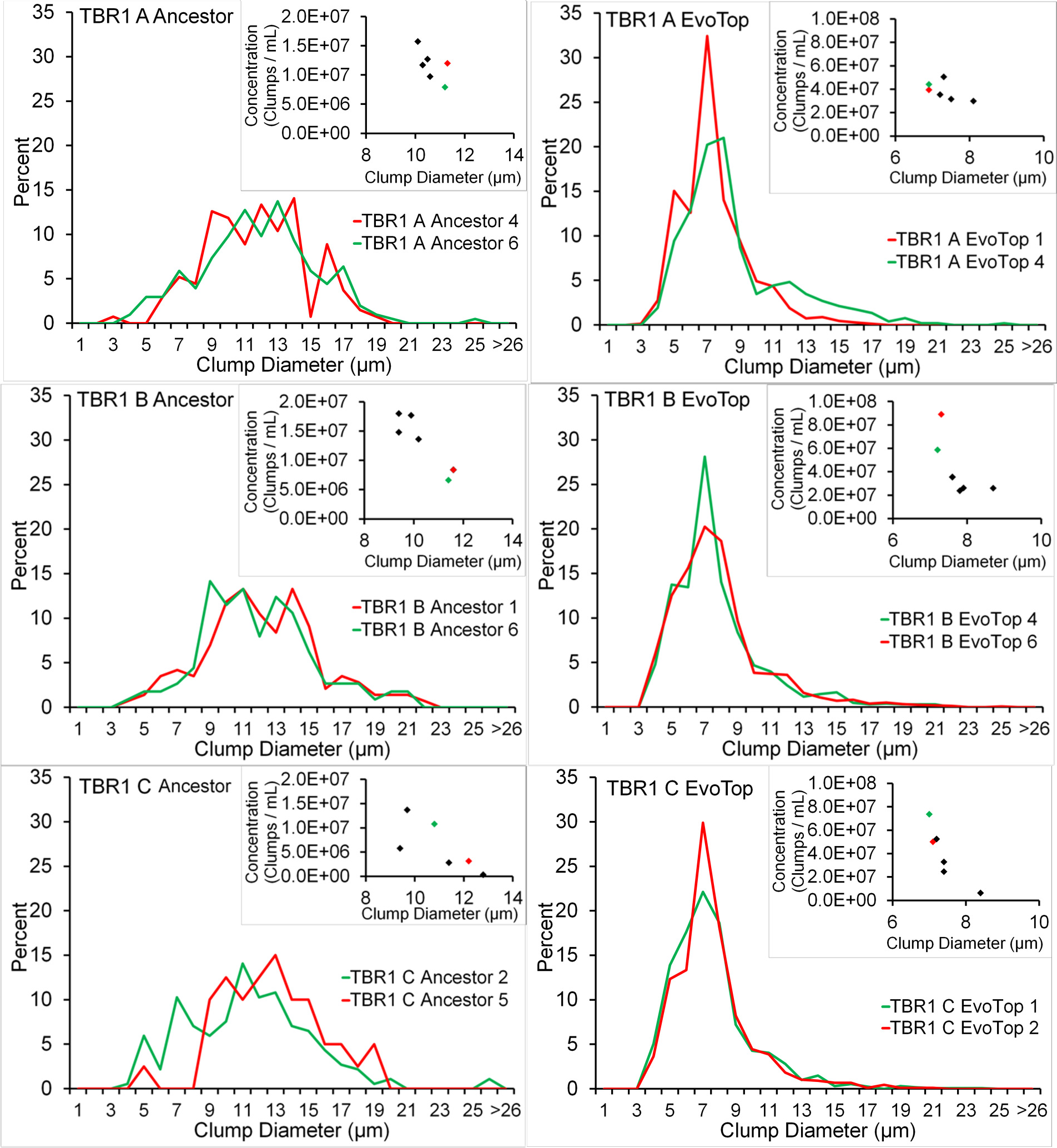
Clump diameter distribution and average size (diamonds) of ancestral (left) and EvoTop (right) isolates. Isolates shown in red were used for the stress test experiments.

**Table S1.** Unique mutations in the TBR1 EvoTop isolates.

| Predicted mutations for the TBR1 A Experimental Isolate |  |  |  |  |  |  |  |
| --- | --- | --- | --- | --- | --- | --- | --- |
| seq id | position | mutation | annotation | gene | description | breseq freq | lofreq* freq |
| gi295413810gbACVY01000005.1 | 346,739 | C→T | G439S (GGC→AGC) | YBR068C_CDS_BAP2 ← | High affinity leucine permease functions as a branched-chain amino acid permease involved in the uptake of leucine isoleucine and valine contains 12 predicted transmembrane domains | 24.13% | 23.70% |
| gi295413810gbACVY01000005.1 | 527,059 | T→G | M1R (ATG→AGG) † | YBR158W_CDS_AMN1 → | Protein required for daughter cell separation multiple mitotic checkpoints and chromosome stability contains 12 degenerate leucine-rich repeat motifs expression is induced by the Mitotic Exit Network MEN | 72.59% | 72.11% |
| gi295413810gbACVY01000005.1 | 527,116 | C→A | S20* (TCA→TAA) | YBR158W_CDS_AMN1 → | Protein required for daughter cell separation multiple mitotic checkpoints and chromosome stability contains 12 degenerate leucine-rich repeat motifs expression is induced by the Mitotic Exit Network MEN | 18.93% | 18.13% |
| gi295413787gbACVY01000028.1 | 56,138 | G→C | S174C (TCC→TGC) | YER088C_CDS_DOT6 ← | Protein involved in rRNA and ribosome biogenesis binds polymerase A and C motif subunit of the RPD3L histone deacetylase complex similar to Tod6p has chromatin specific SANT domain involved in telomeric gene silencing and filamentation | 23.60% | 22.89% |
| gi295413784gbACVY01000031.1 | 47,990 | C→T | E1693K (GAG→AAG) | YLL040C_CDS_VPS13 ← | Protein of unknown function heterooligomeric or homooligomeric complex peripherally associated with membranes homologous to human COH1 involved in sporulation vacuolar protein sorting and protein-Golgi retention | 64.11% | 61.46% |
| Predicted mutations for the TBR1 B Experimental Isolate |  |  |  |  |  |  |  |
| seq id | position | mutation | annotation | gene | description | breseq freq | lofreq* freq |
| gi295413810gbACVY01000005.1 | 528,271 | C→G | T405R (ACG→AGG) | YBR158W_CDS_AMN1 → | Protein required for daughter cell separation multiple mitotic checkpoints and chromosome stability contains 12 degenerate leucine-rich repeat motifs expression is induced by the Mitotic Exit Network MEN | 100.00% | 97.86% |
| gi295413808gbACVY01000007.1 | 431,329 | C→A | Q1442K (CAA→AAA) | YJL005W_CDS_CYR1 → | Adenylate cyclase required for cAMP production and cAMP-dependent protein kinase signaling the cAMP pathway controls a variety of cellular processes including metabolism cell cycle stress response stationary phase and sporulation | 100.00% | 96.89% |
| gi295413805gbACVY01000010.1 | 84,670 | C→A | P71T (CCA→ACA) | YOR255W_CDS_OSW1 → | Protein involved in sporulation required for the construction of the outer spore wall layers required for proper localization of Spo14p | 100.00% | 98.95% |
| gi295413792gbACVY01000023.1 | 63,441 | T→C | intergenic (-405/-334) | YIL023C_CDS_YKE4 ← /<br>→ YIL022W_CDS_TIM44 | Zinc transporter localizes to the ER null mutant is sensitive to calcofluor white leads to zinc accumulation in cytosol ortholog of the mouse KE4 and member of the ZIP ZRT IRT-like Protein family/Essential component of the Translocase of the Inner Mitochondrial membrane TIM23 complex tethers the import motor and regulatory factors PAM complex to the translocation channel Tim23p-Tim17p core complex | 100.00% | 96.43% |
| gi295413769gbACVY01000046.1 | 1,436 | G→T | intergenic (-/-) | -/- | -/- | 100.00% | 100.00% |
| Predicted mutations for the TBR1 C Experimental Isolate |  |  |  |  |  |  |  |
| seq id | position | mutation | annotation | gene | description | breseq freq | lofreq* freq |
| gi295413810gbACVY01000005.1 | 528,545 | A→C | K496N (AAA→AAC) | YBR158W_CDS_AMN1 → | Protein required for daughter cell separation multiple mitotic checkpoints and chromosome stability contains 12 degenerate leucine-rich repeat motifs expression is induced by the Mitotic Exit Network MEN | 100.00% | 96.57% |

Unique to evolved line.
Unique to evolved line's isolate.

## REFERENCES

Acar, M., J. T. Mettetal, and A. van Oudenaarden. 2008. Stochastic switching as a survival strategy in fluctuating environments. Nat Genet 40:471–475.

Aldridge, B. B., M. Fernandez-Suarez, D. Heller, V. Ambravaneswaran, D. Irimia, M. Toner, and S. M. Fortune. 2012. Asymmetry and Aging of Mycobacterial Cells Lead to Variable Growth and Antibiotic Susceptibility. Science 335:100–104.

Andersen, K. S., R. Bojsen, L. G. R. Sorensen, M. W. Nielsen, M. Lisby, A. Folkesson, and B. Regenberg. 2014. Genetic Basis for Saccharomyces cerevisiae Biofilm in Liquid Medium. G3-Genes Genom Genet 4:1671–1680.

Brachmann, C. B., A. Davies, G. J. Cost, E. Caputo, J. C. Li, P. Hieter, and J. D. Boeke. 1998. Designer deletion strains derived from Saccharomyces cerevisiae S288C: a useful set of strains and plasmids for PCR-mediated gene disruption and other applications. Yeast 14:115–132.

Brunke, S., K. Seider, D. Fischer, I. D. Jacobsen, L. Kasper, N. Jablonowski, A. Wartenberg, O. Bader, A. Enache-Angoulvant, M. Schaller, C. d’Enfert, and B. Hube. 2014. One Small Step for a Yeast - Microevolution within Macrophages Renders Candida glabrata Hypervirulent Due to a Single Point Mutation. Plos Pathog 10.

Cap, M., L. Vachova, and Z. Palkova. 2012. Reactive oxygen species in the signaling and adaptation of multicellular microbial communities. Oxid Med Cell Longev 2012:976753.

Chaudhari, R. D., J. D. Stenson, T. W. Overton, and C. R. Thomas. 2012. Effect of bud scars on the mechanical properties of Saccharomyces cerevisiae cell walls. Chem Eng Sci 84:188–196.

Chen, L., J. Noorbakhsh, R. M. Adams, J. Samaniego-Evans, G. Agollah, D. Nevozhay, J. Kuzdzal-Fick, P. Mehta, and G. Balazsi. 2014. Two-Dimensionality of Yeast Colony Expansion Accompanied by Pattern Formation. Plos Comput Biol 10.

Collin, R. and M. P. Miglietta. 2008. Reversing opinions on Dollo’s Law. Trends Ecol Evol 23:602–609.

Deatherage, D. E. and J. E. Barrick. 2014. Identification of Mutations in Laboratory-Evolved Microbes from Next-Generation Sequencing Data Using breseq. Methods Mol Biol 1151:165–188.

Di Talia, S., H. Y. Wang, J. M. Skotheim, A. P. Rosebrock, B. Futcher, and F. R. Cross. 2009. Daughter-Specific Transcription Factors Regulate Cell Size Control in Budding Yeast. Plos Biol 7.

Dollo, L. 1893. Les lois de l’évolution. Bull. Soc. Belge Geol. Pal. Hydr. 7:164–166.

Dowell, R. D., O. Ryan, A. Jansen, D. Cheung, S. Agarwala, T. Danford, D. A. Bernstein, P. A. Rolfe, L. E. Heisler, B. Chin, C. Nislow, G. Giaever, P. C. Phillips, G. R. Fink, D. K. Gifford, and C. Boone. 2010. Genotype to Phenotype: A Complex Problem. Science 328:469–469.

Giaever, G., A. M. Chu, L. Ni, C. Connelly, L. Riles, S. Veronneau, S. Dow, A. Lucau-Danila, K. Anderson, B. Andre, A. P. Arkin, A. Astromoff, M. El Bakkoury, R. Bangham, R. Benito, S. Brachat, S. Campanaro, M. Curtiss, K. Davis, A. Deutschbauer, K. D. Entian, P. Flaherty, F. Foury, D. J. Garfinkel, M. Gerstein, D. Gotte, U. Guldener, J. H. Hegemann, S. Hempel, Z. Herman, D. F. Jaramillo, D. E. Kelly, S. L. Kelly, P. Kotter, D. LaBonte, D. C. Lamb, N. Lan, H. Liang, H. Liao, L. Liu, C. Y. Luo, M. Lussier, R. Mao, P. Menard, S. L. Ooi, J. L. Revuelta, C. J. Roberts, M. Rose, P. Ross-Macdonald, B. Scherens, G. Schimmack, B. Shafer, D. D. Shoemaker, S. Sookhai-Mahadeo, R. K. Storms, J. N. Strathern, G. Valle, M. Voet, G. Volckaert, C. Y. Wang, T. R. Ward, J. Wilhelmy, E. A. Winzeler, Y. H. Yang, G. Yen, E. Youngman, K. X. Yu, H. Bussey, J. D. Boeke, M. Snyder, P. Philippsen, R. W. Davis, and M. Johnston. 2002. Functional profiling of the Saccharomyces cerevisiae genome. Nature 418:387–391.

Gonzalez, C., J. C. J. Ray, M. Manhart, R. M. Adams, D. Nevozhay, A. V. Morozov, and G. Balazsi. 2015. Stress-response balance drives the evolution of a network module and its host genome. Mol Syst Biol 11.

Gregor, T., K. Fujimoto, N. Masaki, and S. Sawai. 2010. The Onset of Collective Behavior in Social Amoebae. Science 328:1021–1025.

Grosberg, R. K. and R. R. Strathmann. 2007. The evolution of multicellularity: A minor major transition? Annu Rev Ecol Evol S 38:621–654.

Harju, S., H. Fedosyuk, and K. R. Peterson. 2004. Rapid isolation of yeast genomic DNA: Bust n’ Grab. Bmc Biotechnol 4.

Klimov, P. B. and B. OConnor. 2013. Is Permanent Parasitism Reversible?-Critical Evidence from Early Evolution of House Dust Mites. Syst Biol 62:411–423.

Koschwanez, J. H., K. R. Foster, and A. W. Murray. 2011. Sucrose Utilization in Budding Yeast as a Model for the Origin of Undifferentiated Multicellularity. Plos Biol 9.

Kuranda, M. J. and P. W. Robbins. 1991. Chitinase Is Required for Cell-Separation during Growth of Saccharomyces-Cerevisiae. J Biol Chem 266:19758–19767.

Kuthan, M., F. Devaux, B. Janderova, I. Slaninova, C. Jacq, and Z. Palkova. 2003. Domestication of wild Saccharomyces cerevisiae is accompanied by changes in gene expression and colony morphology. Mol Microbiol 47:745–754.

Kuzdzal-Fick, J. J., K. R. Foster, D. C. Queller, and J. E. Strassmann. 2007. Exploiting new terrain: an advantage to sociality in the slime mold Dictyostelium discoideum. Behav Ecol 18:433–437.

Levy, S. F., N. Ziv, and M. L. Siegal. 2012. Bet Hedging in Yeast by Heterogeneous, Age-Correlated Expression of a Stress Protectant. Plos Biol 10.

Li, J. R., L. Wang, X. P. Wu, O. Fang, L. W. Wang, C. Q. Lu, S. J. Yang, X. H. Hu, and Z. W. Luo. 2013. Polygenic Molecular Architecture Underlying Non-Sexual Cell Aggregation in Budding Yeast. DNA Res 20:55–66.

Lo, W. S. and A. M. Dranginis. 1998. The cell surface flocculin Flo11 is required for pseudohyphae formation and invasion by Saccharomyces cerevisiae. Mol Biol Cell 9:161–171.

Maynard Smith, J. and E. Szathmáry. 1998. The Major Transitions in Evolution. Oxford University Press.

McCandlish, D. M., P. Shah, and J. B. Plotkin. 2016. Epistasis and the Dynamics of Reversion in Molecular Evolution. Genetics 203:1335–1351.

Paul, G. S. 2002. Dinosaurs of the Air: The Evolution and Loss of Flight in Dinosaurs and Birds. Johns Hopkins University Press.

Pentz, J. T., T. Limberg, N. Beermann, and W. C. Ratcliff. 2015. Predator Escape: An Ecologically Realistic Scenario for the Evolutionary Origins of Multicellularity. Evolution: Education and Outreach 8:13.

Peters, G. 2018. userfriendlyscience: Quantitative analysis made accessible. R package version 0.7.1.

R_Core_Team. 2013. R: A language and environment for statistical computing. R Foundation for Statistical Computing, Vienna, Austria.

Ratcliff, W. C., R. F. Denison, M. Borrello, and M. Travisano. 2012. Experimental evolution of multicellularity. P Natl Acad Sci USA 109:1595–1600.

Ratcliff, W. C., J. D. Fankhauser, D. W. Rogers, D. Greig, and M. Travisano. 2015. Origins of multicellular evolvability in snowflake yeast. Nat Commun 6.

Reynolds, T. B. and G. R. Fink. 2001. Bakers’ yeast, a model for fungal biofilm formation. Science 291:878–881.

Saxer, G., M. D. Krepps, E. D. Merkley, C. Ansong, B. L. D. Kaiser, M. T. Valovska, N. Ristic, P. T. Yeh, V. P. Prakash, O. P. Leiser, L. Nakhleh, H. S. Gibbons, H. W. Kreuzer, and Y. Shamoo. 2014. Mutations in Global Regulators Lead to Metabolic Selection during Adaptation to Complex Environments. Plos Genet 10.

Shoval, O., H. Sheftel, G. Shinar, Y. Hart, O. Ramote, A. Mayo, E. Dekel, K. Kavanagh, and U. Alon. 2012. Evolutionary Trade-Offs, Pareto Optimality, and the Geometry of Phenotype Space. Science 336:1157–1160.

Smith, J., D. C. Queller, and J. E. Strassmann. 2014. Fruiting bodies of the social amoeba Dictyostelium discoideum increase spore transport by Drosophila. Bmc Evol Biol 14.

Smukalla, S., M. Caldara, N. Pochet, A. Beauvais, S. Guadagnini, C. Yan, M. D. Vinces, A. Jansen, M. C. Prevost, J. P. Latge, G. R. Fink, K. R. Foster, and K. J. Verstrepen. 2008. FLO1 Is a Variable Green Beard Gene that Drives Biofilm-like Cooperation in Budding Yeast. Cell 135:726–737.

Soylemez, O. and F. A. Kondrashov. 2012. Estimating the Rate of Irreversibility in Protein Evolution. Genome Biol Evol 4:1213–1222.

Wang, Y. C., T. Shirogane, D. Liu, J. W. Harper, and S. J. Elledge. 2003. Exit from exit: Resetting the cell cycle through Amn1 inhibition of G protein signaling. Cell 112:697–709.

Wei, W., J. H. McCusker, R. W. Hyman, T. Jones, Y. Ning, Z. Cao, Z. Gu, D. Bruno, M. Miranda, M. Nguyen, J. Wilhelmy, C. Komp, R. Tamse, X. Wang, P. Jia, P. Luedi, P. J. Oefner, L. David, F. S. Dietrich, Y. Li, R. W. Davis, and L. M. Steinmetz. 2007. Genome sequencing and comparative analysis of Saccharomyces cerevisiae strain YJM789. P Natl Acad Sci USA 104:12825–12830.

Wilm, A., P. P. K. Aw, D. Bertrand, G. H. T. Yeo, S. H. Ong, C. H. Wong, C. C. Khor, R. Petric, M. L. Hibberd, and N. Nagarajan. 2012. LoFreq: a sequence-quality aware, ultra-sensitive variant caller for uncovering cell-population heterogeneity from high-throughput sequencing datasets. Nucleic Acids Res 40:11189–11201.

Wloch-Salamon, D. M., M. Plech, and J. Majewska. 2013. Generation of stable, non-aggregating Saccharomyces cerevisiae wild isolates. Acta Biochim Pol 60:657–660.

Yvert, G., R. B. Brem, J. Whittle, J. M. Akey, E. Foss, E. N. Smith, R. Mackelprang, and L. Kruglyak. 2003. Trans-acting regulatory variation in Saccharomyces cerevisiae and the role of transcription factors. Nat Genet 35:57–64.

Zara, G., S. Zara, C. Pinna, S. Marceddu, and M. Budroni. 2009. FLO11 gene length and transcriptional level affect biofilm-forming ability of wild flor strains of Saccharomyces cerevisiae. MicrobiolSgm 155:3838–3846.

